# Inhibition of polo-like kinase 1 (PLK1) facilitates reactivation of gamma-herpesviruses in B-cell lymphomas and their elimination

**DOI:** 10.1101/2020.10.08.330548

**Authors:** Ayan Biswas, Guillaume N. Fiches, Dawei Zhou, Zhenyu Wu, Xuefeng Liu, Qin Ma, Weiqiang Zhao, Jian Zhu, Netty G. Santoso

## Abstract

Both Kaposi’s sarcoma-associated herpesvirus (KSHV) and Epstein-Barr virus (EBV) establish the persistent, life-long infection primarily at the latent status, and contribute to certain types of tumors, including B cell lymphomas, especially in immuno-compromised individuals, such as people living with HIV (PLWH). Lytic reactivation of these viruses can be employed to kill tumor cells harboring latently infected viral episomes, through the viral cytopathic effects and the subsequent antiviral immune responses. In this study, we identified that polo-like kinase 1 (PLK1) is induced by KSHV *de novo* infection as well as lytic switch from KSHV latency. We further demonstrated that PLK1 depletion or inhibition facilitates KSHV reactivation and promotes cell death of KSHV-reactivated lymphoma cells. Mechanistically, PLK1 regulates Myc that is critical to both maintenance of KSHV latency and support of cell survival, and preferentially affects the level of H3K27me3 inactive mark both globally and at certain loci of KSHV viral episomes. Lastly, we recognized that PLK1 inhibition synergizes with STAT3 inhibition to efficiently induce KSHV reactivation. Furthermore, we also confirmed that PLK1 depletion or inhibition yields the similar effect on EBV reactivation and cell death of EBV-reactivated lymphoma cells. Additionally, we noticed the PLK1 in B cells is elevated in the context of HIV infection and by HIV Nef protein, which would favor KSHV/EBV latency. Above all, our findings illustrated that PLK1 is a novel host target that can be inhibited to benefit the viral oncolysis to eliminate KSHV/EBV-infected lymphoma cells, particularly for PLWH.

## Introduction

Kaposi’s sarcoma-associated herpesvirus (KSHV), is an etiological agent of Kaposi’s sarcoma (KS) – a common AIDS-associated malignancy^1^, as well as two lymphoproliferative diseases, namely primary effusion lymphoma (PEL) and multicentric Castleman’s disease (CAD)^2,3^. KSHV has a diverse range of *in vivo* and *in vitro* cell tropism, but CD19+ B cells appear to be the primary cell type of KSHV persistent infection^4,5^. As similar to other herpesviruses, KSHV infection also includes latent and lytic phases ^6,7^. Following acute infection, KSHV establishes latency in most of immunocompetent individuals, primarily in B cells^7^. Such KSHV latently infected B cells constitute a major source of KSHV viral reservoirs to maintain KSHV genomes and propagate new KSHV viruses. At the latent phase, only a limited subset of KSHV latent genes is expressed, while most of lytic genes is silenced. Viral latency supports the maintenance of viral episomes, but meanwhile escapes from the host immune surveillance. Latent KSHV can be reactivated in response to certain stimuli, and expression of viral lytic genes is turned on to produce new virions during viral lytic cycle. Epstein-barr virus (EBV) belongs to the same human γ-herpesvirus family as KSHV, and also primarily infect B cells in a latent phage that can be reserved to lytic replication. It is interesting to note that nearly 70% of PEL cell lines are co-infected with EBV. Studies have demonstrated that EBV co-infected PEL cell lines are more tumorigenic compared to the EBV negative ones^8,9^.

Oncolytic viruses have been recently engineered as novel anticancer agents and shown to increase the therapeutic promise^10,11^. Similarly, lytic reactivation of intrinsic latent viruses, such as human endogenous retroviruses (HERVs), in tumor cells can also lead to oncolysis and be applied as an anticancer therapy^12^. The similar approach has been attempted for elimination of KSHV-infected tumor cells through viral oncolysis. Agonists or antagonists modulating certain cell signaling have been employed to reactivate latent KSHV, which has been demonstrated to eliminate KSHV-infected tumor cells^13,14^. This is mostly due to the virus-induced cytopathic effect that promotes the death of virus-reactivated cell. Although these findings are promising, such approaches are still at the infant stage and have received only limited investigation so far. Therefore, we are interested in understanding the common cell mechanisms associated with not only reactivation of latent KSHV and EBV but also cell survival of KSHV/EBV-reactivated tumor cells, which can be employed as a therapeutic strategy to eliminate the tumor cells harboring these oncoviruses.

PLK1 is the member of the serine/threonine polo-like kinase (PLK) family (PLK 1-4), which are key players of multiple cellular functions, including cell cycle progression, cellular stress response, and innate immune signaling^15,16,17^. PLK1 expression is often elevated in various human cancers, and links to tumor aggressiveness and poor clinical prognosis. In particular, as a relatively new mechanism PLK1 plays an important role in promoting cell survival through stabilization of Myc protein in tumors, including B-cell lymphomas^18,19^. Recently, we have reported that PLK1 protein is elevated due to HIV viral infection in CD4^+^ T cells and contributes to cell survival^20^. However, there are currently no reports regarding to PLK1’s role in regulation of viral infection and oncogenesis for human gamma-herpesviruses. In this study, we identified that PLK1 expression is induced by KSHV viral infection and plays a critical role in maintaining latent infection of KSHV in B lymphoma cells as well as supporting their cell survival status via Myc protein. We further showed that specific inhibition of PLK1 facilitates viral lytic reactivation of KSHV and elimination of B lymphoma cells harboring KSHV viral reservoirs. We also observed the similar functions of PLK1 in EBV-infected B lymphoma cells. PLK1 inhibitors have already demonstrated the promising results to inhibit cell survival of tumor cells *vs* non-tumor cells^21,22^, and our studies suggest that they can be further considered for viral oncolysis approach to treat KSHV/EBV-positive B lymphomas particularly for PLWH, given that we identified that PLK1 expression in B cells is also induced in context of HIV infection.

## Results

### PLK1 is induced by KSHV infection and required for KSHV latency

We first determined the impact of KSHV infection on PLK1 expression. SLK cells, a renal carcinoma cell line with epithelial-cell origin, were spinoculated with KSHV.BAC16 viruses. KSHV infection rate was quantified by measuring the GFP expression from KSHV viral genomes in the infected cells. Fluorescence imaging showed that KSHV *de novo* infection significantly induces the expression of PLK1 protein by the immunofluorescence assays (IFAs) in SLK cells at 24hpi **(Fig 1A)**. We also evaluated the PLK1 protein expression in the scenario of KSHV lytic switch from latency. iSLK.BAC16 cells^23^ were treated with either Dox (doxycycline) to induce KSHV RTA expression and consequently lytic reactivation, or vehicle control to keep KSHV at latency. Fluorescence imaging confirms that KSHV lytic reactivation also significantly induces the expression of PLK1 protein by IFAs in iSLK.BAC16 cells **(Fig 1B)**. Dox treatment has no effect on PLK1 protein expression in SLK cells.

**Figure 1.**
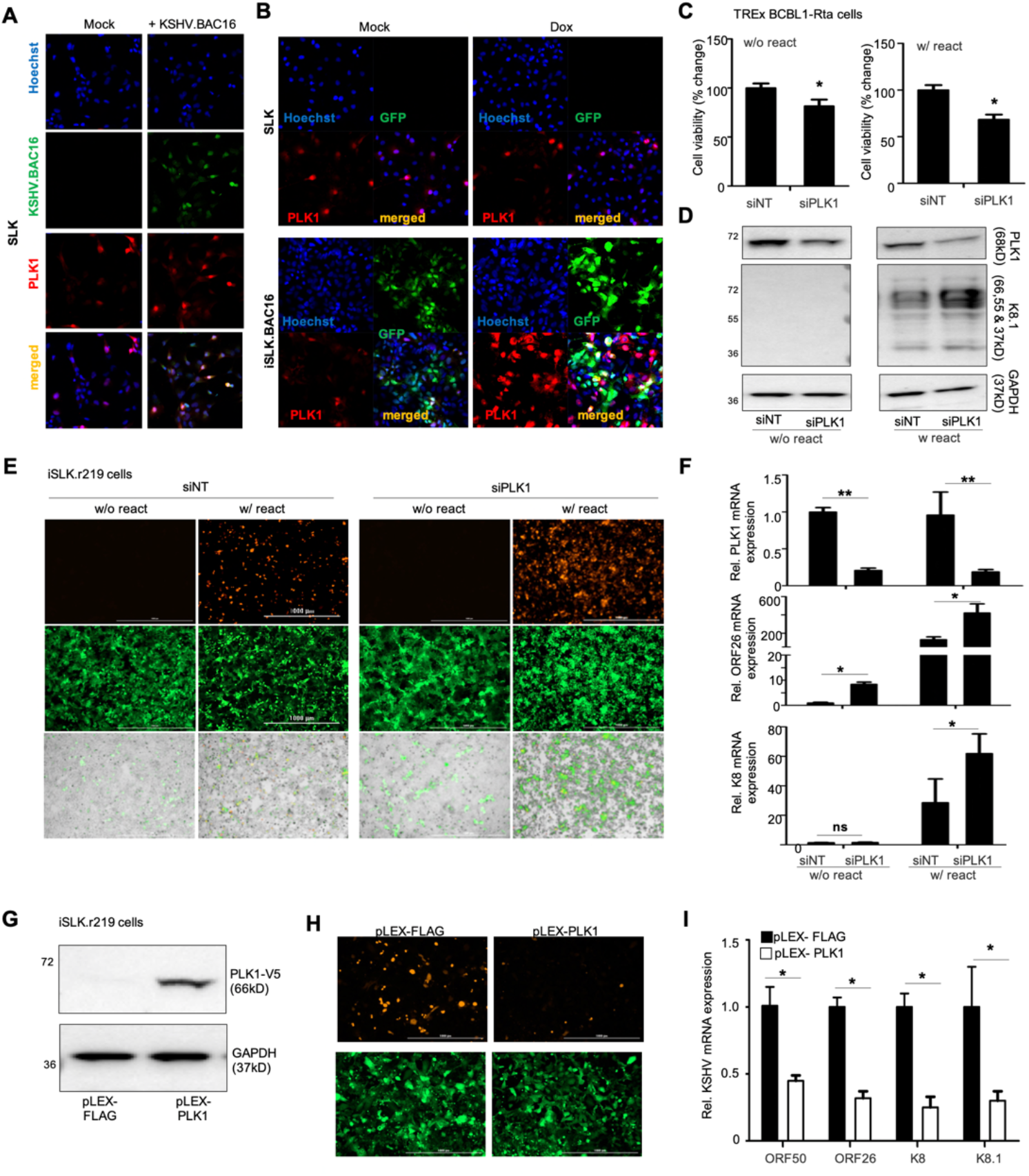
PLK1 plays a role in maintenance of KSHV latency. (A) SLK cells were *de novo* infected with KSHV.BAC16 viruses that were harvested from supernatant of iSLK.BAC16 cell treated with doxycycline (Dox). KSHV-infected SLK cells were fixed, stained with an anti-PLK1 primary antibody and an Alexa Fluor™ 647 labeled secondary antibody, followed by the confocal fluorescence imaging. Nuclei were stained with Hoechst (blue), and GFP indicated the cells were infected with KSHV.BAC16 viruses (green). (B) SLK and iSLK.BAC16 cells were treated with Dox or mock, followed by the immunostaining of PLK1 and confocal fluorescence imaging. Images were one representative from three independent experiments. (C) TREx BCBL1-Rta cells were transiently transfected with the indicated siRNAs (siNT or siPLK1) and treated with Dox or mock to induce KSHV reactivation, followed by the cell viability analysis. (D) Protein level of PLK1 and KSHV lytic gene K8.1 in above cells (C) was measured by immunoblotting. GAPDH was used as a loading control. (E) iSLK.r219 cells were transiently transfected with indicated siRNAs and treated with Dox or mock to induce KSHV reactivation. GFP (constitutively expressed from EF-1alpha promoter within KSHV genome) and RFP (expressed from KSHV PAN lytic promoter) signals in cells were visualized by fluorescence microscopy. (F) mRNA level of PLK1 and KSHV lytic genes (ORF26, K8) in above cells (E) were measured by RT-qPCR and normalized to GAPDH. (G) iSLK.r219 cells were transiently transfected with pLEX-FLAG or pLEX-PLK1 vector, and treated with Dox to induce KSHV reactivation. Expression of V5-PLK1 protein was confirmed by immunoblotting. (H) GFP and RFP signals in above cells (G) were visualized by fluorescence microscopy. (I) mRNA level of KSHV lytic genes (ORF50, ORF26, K8, K8.1) in above cells (G) were measured by RT-qPCR and normalized to GAPDH. Results were calculated from n=3 independent experiments and presented as mean ± SD (* p<0.05, ** p<0.01; two-tailed paired Student t-test).

To investigate the role of PLK1 in regulating KSHV infection, we depleted the expression of endogenous PLK1 in TREx BCBL-Rta cells using its siRNA (siPLK1) through electroporation, while non-target siRNA (siNT) was used as a control. PLK1 depletion led to the slightly more reduction of cell viability, especially in Dox-treated, i.e. KSHV-reactivated, cells **(Fig 1C)**. siPLK1 knockdown efficiency was confirmed by immunoblotting of PLK1 protein in these cells **(Fig 1D)**. Meanwhile, we noticed that PLK1 depletion significantly increases the KSHV lytic reactivation specifically in Dox-treated cells but not the un-treated ones through measurement of K8.1 lytic protein by immunoblotting (**Fig 1D**). We observed the similar effect of siRNA-mediated PLK1 depletion (**Fig 1F**) on KSHV lytic switch from latency in iSLK.r219 cells through measurement of RFP expression from KSHV lytic PAN promoter by fluorescent imaging (**Fig 1E**) or KSHV lytic gene expression (ORF26, K8) by RT-qPCR (**Fig 1F**). These results indicated that presence of PLK1 protein is necessary but not sufficient for maintenance of KSHV latency. Alternatively, we also confirmed such a function of PLK1 by gain-of-function analysis. A pLEX vector expressing V5-tagged PLK1 (pLEX-PLK1) or FLAG peptide (pLEX-FLAG) was transiently transfected in iSLK.r219 cells, and its expression was confirmed by protein immunoblotting (**Fig 1G**) or RT-qPCR (**Fig S1**). Exogenous expression of PLK1 led to the significant counteraction of Dox-induced KSHV lytic reactivation in these cells through measurement of RFP by fluorescent imaging (**Fig 1H**) or KSHV lytic genes expression (ORF50, ORF26, K8, K8.1) by RT-qPCR (**Fig 1I**).

### Inhibition of PLK1 facilitates the KSHV lytic switch from latency

Our recent studies demonstrated that inhibition of PLK1 enables the reactivation of latent HIV^20^. Since we found that PLK1 depletion facilitates KSHV lytic reactivation, we further evaluated whether inhibition of PLK1 yields the similar effect. We used a PLK1-specific inhibitor, SBE 13 HCl (SBE), which targets PLK1 inactive kinase conformation^24^. Consistent with the results of PLK1 depletion, SBE treatment led to the further increase of Dox-induced KSHV lytic reactivation in TREx BCBL1-Rta cells in a dose dependent manner through measurement of KSHV K8.1 lytic protein by immunoblotting (**Fig 2A**) or lytic gene expression (ORF50, K8, K8.1) by RT-qPCR (**Fig 2B**). However, treatment of SBE alone is not sufficient to induce any KSHV lytic reactivation in TREx BCBL1-Rta cells (data not shown). We also tested SBE in the parental BCBL1 cells, a PEL cell line latently infected with KSHV. BCBL1 cells were treated with 12-O-tetra-decanoylphorbol-13-acetate (TPA) and sodium butyrate (TPA/NaB) or mock treated in the presence or absence of SBE. SBE alone had certain weak effect to induce KSHV lytic reactivation, but SBE further dramatically enhanced the KSHV latency-reversing potency of TPA/NaB (**Fig 2C**). Likewise, SBE treatment alone failed to reactivate latent KSHV but synergized with Dox to efficiently induce KSHV lytic reactivation in iSLK.r219 cells as well through measurement of RFP by fluorescent imaging (**Fig 2D**) or KSHV ORF45 lytic protein by immunoblotting (**Fig 2E**), although SBE treatment didn’t cause any obvious cytotoxicity in these cells (**Fig S2A**).

**Figure 2.**
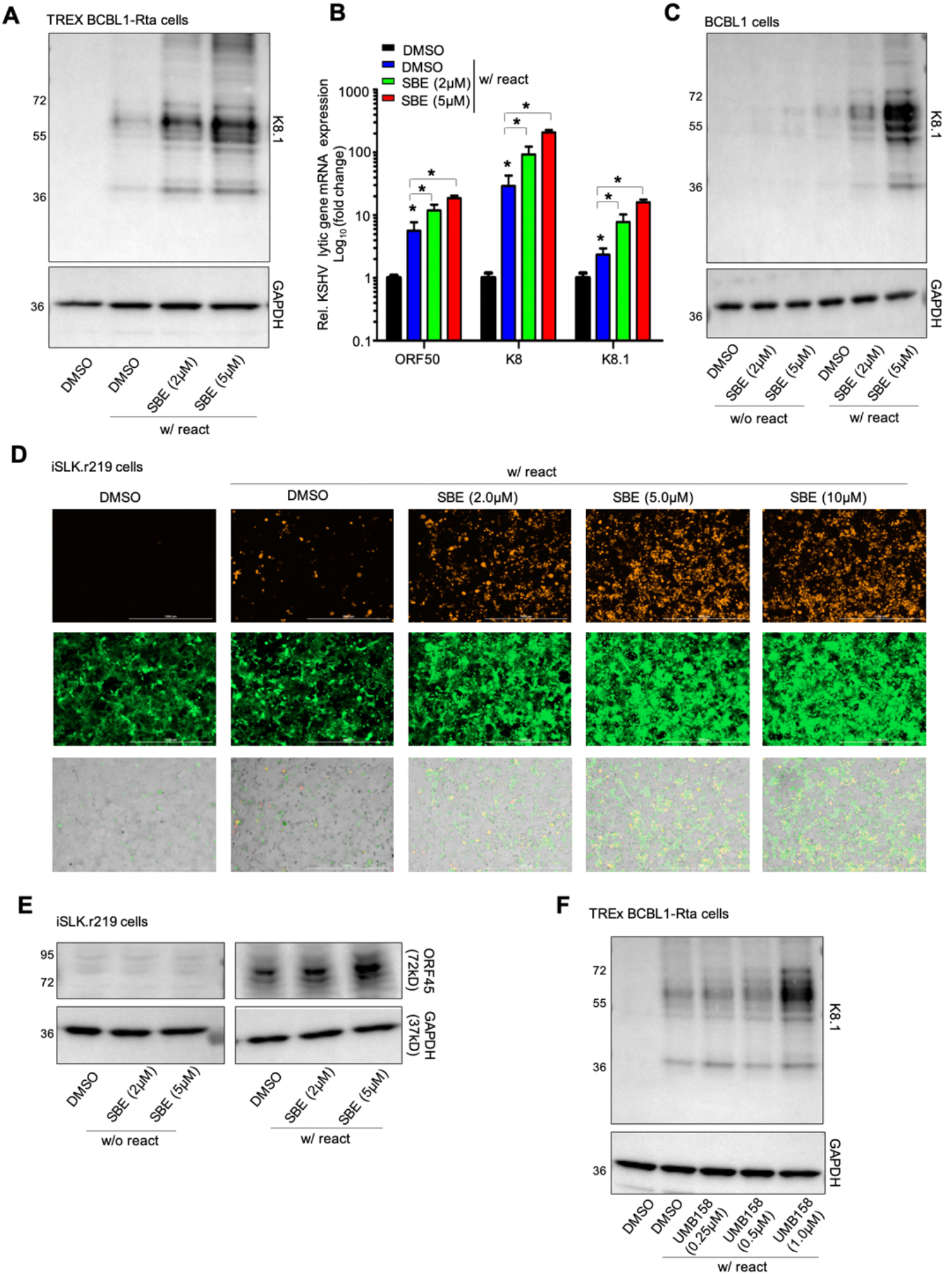
PLK1 inhibition facilitates lytic switch of latent KSHV. (A) TREx BCBL1-Rta cells were treated with the PLK1-specific inhibitor, SBE 13 HCl (SBE), and Dox to induce KSHV reactivation. Protein level of KSHV K8.1 was measured by immunoblotting. GAPDH was used as a loading control. (B) mRNA level of KSHV lytic genes (ORF50, K8, K8.1) in above cells (A) were measured by RT-qPCR and normalized to GAPDH. (C) BCBL1 cells were treated with SBE or DMSO, in the presence or absence of TPA (20ng/ml) + sodium butyrate (1mM) (TPA/NaB) to induce KSHV reactivation. Protein level of KSHV K8.1 was measured by immunoblotting. GAPDH was used as a loading control. (D) iSLK.r219 cells were treated with SBE or DMSO, and Dox to induced KSHV reactivation. GFP and RFP signals in these cells were visualized by fluorescence microscopy. (E) Protein level of KSHV lytic gene ORF45 in iSLK.r219 cells +/- SBE and +/- Dox was measured by immunoblotting. GAPDH was used as a loading control. (F) TREx BCBL1-Rta cells were treated with the PLK1/BET dual inhibitor, UMB-158, and Dox to induce KSHV reactivation. Protein level of KSHV K8.1 was measured by immunoblotting. GAPDH was used as a loading control. Results were calculated from n=3 independent experiments and presented as mean ± SD (* p<0.05; two-tailed paired Student t-test).

Recently, PLK1-BET dual inhibitors have been developed to target both PLK1 and BRD4, two major protein targets of the anticancer therapies^25,26,27^. We tested one of such PLK1-BET dual inhibitors, UMB-158, which possesses the novel scaffold (UMB series)^28^ and has been lately shown to reactivate latent HIV by our group^20^. Indeed, UMB-158 treatment also further enhanced the Dox-induced KSHV lytic reactivation in TREx BCBL1-Rta cells in a dose dependent manner (**Fig 2F**), but UMB-158 alone failed to reactivate latent KSHV in these cells (data not shown), similar as SBE. We also tested two other PLK1-BET dual inhibitors, BI-6727 and BI-2536, which have been previously reported to reactivate latent HIV^29^. Both of these two PLK1-BET dual inhibitors enhanced Dox-induced KSHV lytic reactivation **(Fig S2B)**.

### Inhibition of PLK1 promotes the cell death of KSHV-reactivated tumor cells

It has been well reported that PLK1 supports cell survival^30,31,20^. Thus, we wondered whether PLK1 inhibition generates a synergistic “killing” effect with viral cytopathic effect caused by KSHV reactivation to further promote the cell death of KSHV-reactivated tumor cells. Indeed, we identified that SBE treatment causes more cell death of TREx BCBL1-Rta cells undergoing Dox-induced KSHV lytic reactivation in comparison with mock treatment measured by the LIVE/DEAD assay (**Fig 3A**). In parallel, we also observed that SBE treatment leads to more reduction of cell viability in Dox-treated TREx BCBL1-Rta cells in comparison with mock treatment measured by the ATP assay although cell viability of these cells in both cases is lower than BJAB cells, a KSHV/EBV-negative B lymphoma cell line, which undergoes the same treatments (**Fig S3A**). Similar effect of SBE was observed in TPA/NaB-treated vs un-treated BCBL1 cells except that cell viability of BCBL1 cells without TPA/NaB induction is similar to that of BJAB cells (**Fig S3B**). UMB-158 showed the comparable effect as SBE to promote more cell death of TREx BCBL1-Rta cells treated with Dox to reactivate latent KSHV comparing to mock treatment (**Fig 3B**). Consistently, UMB-158 treatment leads to more reduction of cell viability in Dox-treated TREx BCBL1-Rta cells in comparison with mock treatment measured by the ATP assay (**Fig S3C**). PLK1-BET dual inhibitors, BI-6727 and BI-2536, also slightly reduced cell viability of Dox-treated TREx BCBL1-Rta cells at the higher dose (**Fig S3D**).

**Figure 3.**
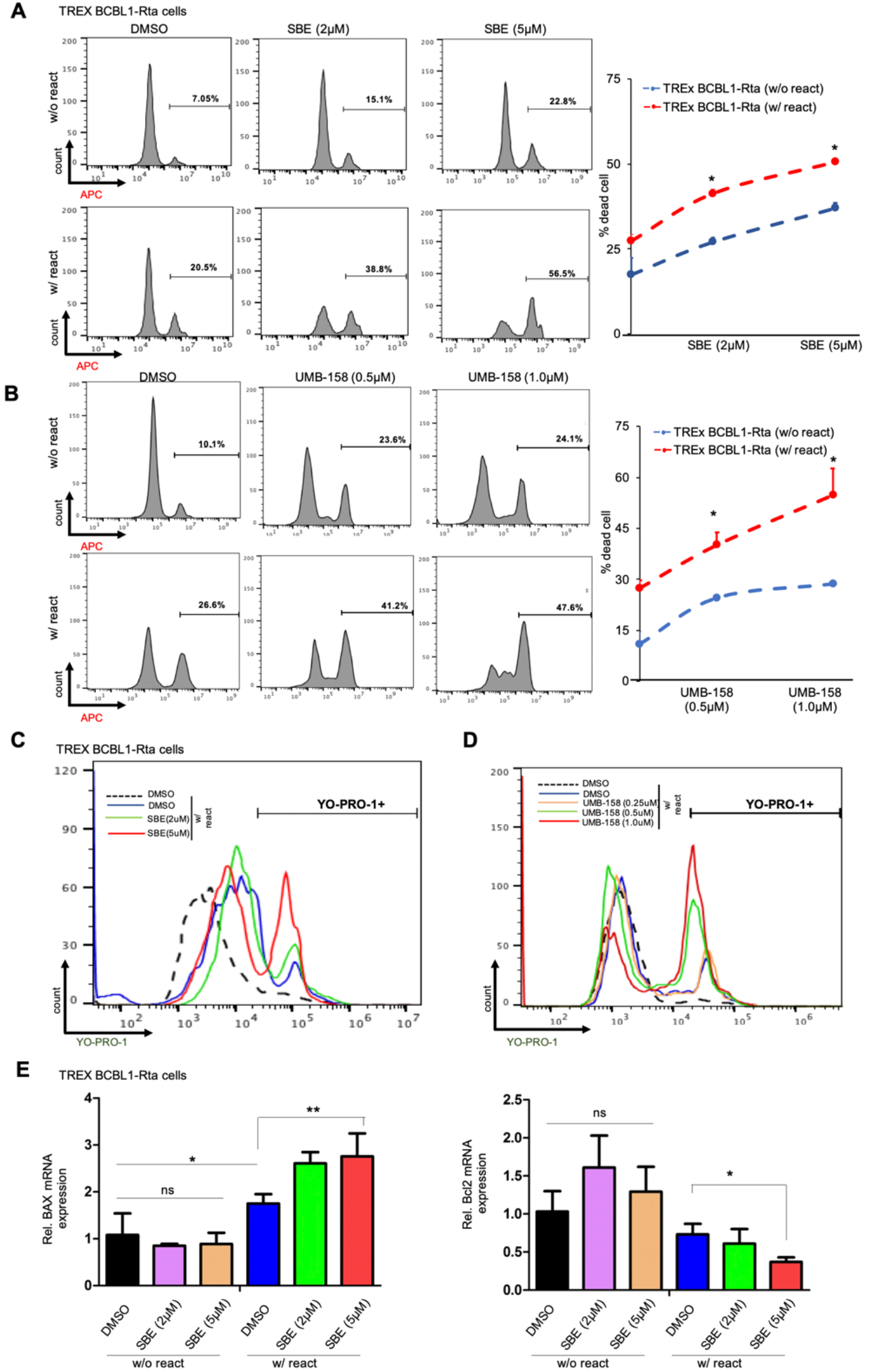
PLK1 inhibition promotes death of KSHV-reactivated cells. (A, B) TREx BCBL1-Rta cells were treated with SBE (A), UMB-158 (B), or DMSO, in the presence or absence of Dox to induce KSHV reactivation. Percentage of dead cells was measured by LIVE/DEAD staining, followed by flow cytometry analysis. (C, D) TREx BCBL1-Rta cells were treated with SBE (C), UMB-158 (D), or DMSO, and induced with Dox. Percentage of apoptotic cells was measured by YO-PRO-1 staining, followed by flow cytometry analysis. (E) mRNA level of apoptosis-related genes, BAX and Bcl2, in TREx BCBL1-Rta cells +/- SBE and +/- Dox were measured by RT-qPCR and normalized to GAPDH. Results were calculated from n=3 independent experiments and presented as mean ± SD (ns: not significant, * p<0.05, ** p<0.01; two-tailed paired Student t-test).

To determine whether SBE and UMB-158 promotes cell death of KSHV-reactivated tumor cells via cell apoptosis, we relied on YO-PRO-1 and propidium iodide (PI) dual staining assays. Apoptotic cells become permeant to green-fluorescent YO-PRO-1, while permissibility of red-fluorescent PI indicates necrotic cell death. Indeed, treatment of SBE (**Fig 3C, S3E**) or UMB-158 (**Fig 3D, S3F**) led to more cell death as well as apoptosis of KSHV-reactivated TREx BCBL1-Rta cells compared to DMSO by measuring the fluorescence intensity at PE and FITC channels respecitively. Furthermore, we also measured the effect of SBE on gene expression of the pro-apoptotic gene BAX^32^ and the anti-apoptotic gene BCL2^33^. SBE led to the upregulation of BAX gene but the downregulation of Bcl2 gene in TREx BCBL1-Rta cells, especially at the higher dose (5 uM), only in the scenario of KSHV reactivation (Dox) but not mock treatment (**Fig 3E**).

### Inhibition of PLK1 destabilizes c-Myc and affects histone methylation

The proto-oncogene myc encodes the c-Myc, a transcription factor, which regulates cellular growth, proliferation, differentiation, and apoptosis^34,18^. It is known that PLK1 regulates c-Myc activities^19,18^, while c-Myc suppresses KSHV lytic reactivation^35^. To evaluate the role of c-Myc in mediating the effect of PLK1 inhibition on promoting KSHV lytic reactivation and cell death of KSHV-reactivated cells, we measured the protein level of c-Myc in iSLK.BAC16 and iSLK.r219 cells treated with SBE, which led to the reduction of c-Myc protein in these cells in a dose-dependent manner **(Fig 4A,B)**. We also tried to measure the c-Myc protein in TREx BCBL1-Rta cells with the similar approach. However, the exogenous Rta protein is Myc-tagged in these cells, which interferes with the immunoblotting of endogenous c-Myc (data not shown). In parallel, we also showed that SBE treatment indeed downregulates the expression of certain Myc-dependent cell-cycle genes, including cyclin A and E (**Fig S4A**), in Dox-treated, KSHV-reactivated TREx BCBL1-Rta cells, further reducing their cell survival.

**Figure 4.**
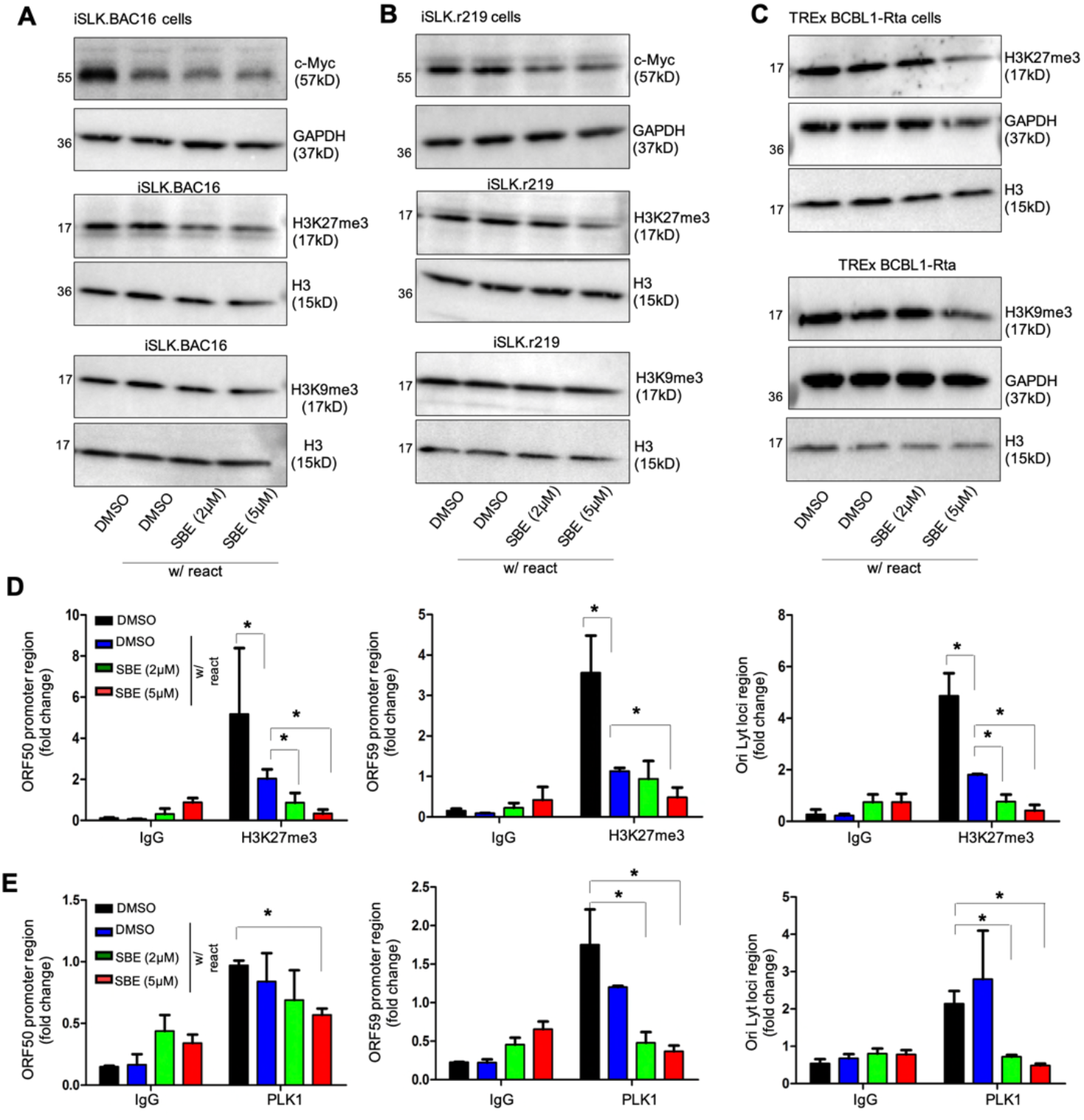
PLK1 inhibition destabilizes c-Myc and reduces H3K27me3. (A-C) Protein level of c-Myc, H3K27me3, H3K9me3, and H3 in iSLK.BAC16 (A), iSLK.r219 (B), or TREx BCBL1-Rta (C) cells was measured by immunoblotting. These cells were treated with SBE or DMSO, and Dox to induce KSHV reactivation. GAPDH was used as a loading control. (D, E) TREx BCBL1-Rta cells were treated with SBE or DMSO, and Dox to induce KSHV reactivation. Cell lysates were prepared and subjected to ChIP assay by using antibodies against H3K27me3, PLK1, or a mouse IgG as a negative control. Precipitated DNA samples were further analyzed by qPCR by using primers targeting promoter region of KSHV lytic genes (ORF50, ORF59) and OriLyt, and normalized to IgG control. Results were calculated from n=3 independent experiments and presented as mean ± SD (* p<0.05; two-tailed paired Student t-test).

It is known that c-Myc modulates the activities of histone lysine methyltransferases^36^. Meanwhile, it has been reported that two inactive histone marks, H3K27me3 and H3K9me3, are involved in maintenance of KSHV latency^37,38^. Therefore, we speculated that inhibition of PLK1 further modulates the above histone marks via destabilization of c-Myc. Our results showed that SBE treatment results in the reduction of overall H3K27me3 but not H3K9me3 histone mark in both iSLK.BAC16 and iSLK.r219 cells (**Fig 4A,B**). In SBE-treated TREx BCBL1-Rta cells, overall H3K27me3 also significantly reduced at the higher dose of SBE (5 uM), while it only caused a mild reduction of H3K9me3 (**Fig 4C**). Furthermore, we determined the effect of SBE on the local level of above histone marks (H3K27me3, H3K9me3) as well as PLK1 at certain loci of KSHV episomes, including promoter region of KSHV lytic genes (ORF50, ORF59) as well as the lytic origin of KSHV DNA replication (OriLyt) by ChIP-PCR. Our results showed that SBE treatment results in the significant reduction of H3K27me3 (**Fig 4D**) and PLK1 (**Fig 4E**) at all above tested loci of KSHV episomes, but not H3K9me3 **(Fig S4B)**.

### Inhibition of PLK1 and STAT3 leads to efficient KSHV lytic switch from latency

As we described previously, inhibition of PLK1 alone is not sufficient to induce efficient KSHV lytic reactivation but rather further promotes it in combination with other latency-reversing agents (LRAs). It has been reported that STAT3 inhibition is sufficient to induce KSHV lytic reactivation^39^. Several STAT3 inhibitors have shown the promising anticancer potency and are currently under evaluation in clinical trials^40,41,42^. We further tested whether the combinatory inhibition of PLK1 and STAT3 induces the efficient KSHV lytic reactivation in BCBL1 cells. Our studies involved two STAT3 inhibitors, cryptotanshinone (CRYP) and stattic, which have been demonstrated to reactivate latent KSHV^43,44^. BCBL1 cells were treated with CRYP (5, 10uM) or stattic (0.25, 0.5uM) alone or in combination with SBE at the lower dose (2uM). CRYP had a moderate KSHV latency-reversing potency, but its combination with SBE enhanced it in BCBL1 cells through measurement of KSHV lytic gene expression (ORF50, K8) by RT-qPCR (**Fig 5A**) as well as K8.1 lytic protein level by immunoblotting (**Fig 5B**). This correlated with the reduced cell viability by the ATP assay (**Fig 5C**). More strikingly, combination of stattic with SBE created a strong KSHV latency-reversing effect superior to the positive control (TPA/NaB treatment) in BCBL1 cells (**Fig 5D,E**), which also correlated with the reduced cell viability (**Fig 5F**). In addition, combination of CRYP or static with SBE even further enhanced the TPA/NaB-induced KSHV lytic reactivation in BCBL1 cells **(Fig S5)**.

**Figure 5.**
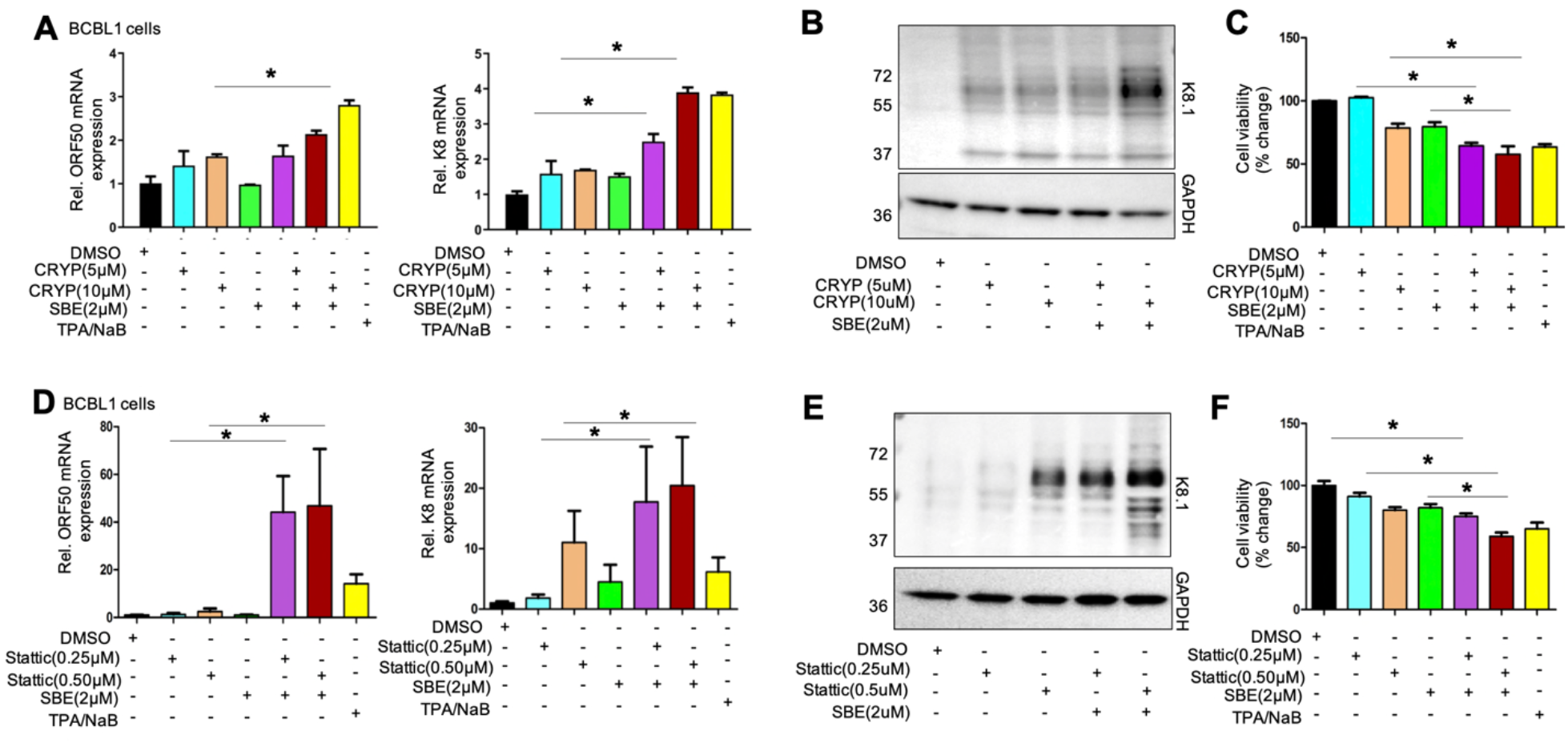
Inhibition of PLK1 and STAT3 efficiently induces KSHV lytic reactivation. (A, D) BCBL1 cells were treated with cryptotanshinone (CRYP, A) or stattic (D) in the presence or absence of SBE. TPA/NaB treatment was used as a positive control to reactivate latent KSHV. mRNA level of KSHV lytic genes (ORF50, K8) in these cells were measured by RT-qPCR and normalized to GAPDH. (B, E) Protein level of KSHV K8.1 in above cells (A, D) was measured by immunoblotting. GAPDH was used as a loading control. (C, F) Cell viability of above cells (A, D) was measured by ATP-based assays and normalized to DMSO. Results were calculated from n=3 independent experiments and presented as mean ± SD (* p<0.05; two-tailed paired Student t-test).

### Inhibition of PLK1 promotes EBV lytic switch and cell death of EBV+ tumor cells

As KSHV shares certain similarities as EBV in host regulation of viral latency and lytic replication^45^, we determined whether PLK1 also regulates EBV lytic switch from latency. We depleted the endogenous PLK1 protein in Akata-BX1 cells by transient transfection of its shRNA (shPLK1) or non-target shRNA (shNT) using TurboFect™ transfection reagent, whose knockdown efficiency was confirmed by immunoblotting **(Fig 6A)**. Such PLK1 knockdown led to the increase of EBV ZEBRA protein (**Fig 6A**) as well as the expression of a series of EBV lytic genes measured by RT-qPCR (**Fig 6B**) in Akata-BX1 cells treated with human IgG (hIgG) to reactivate latent EBV. Furthermore, treatment of SBE (**Fig 6C**) or UMB-158 (**Fig 6D**) indeed further promoted the cell death and reduced the cell viability (**Fig S6A**) of hIgG-treated, EBV-reactivated Akata-BX1 cells, which correlatedly enhanced EBV lytic reactivation through measurement of EBV ZEBRA protein (**Fig 6E,F**). At last, we also evaluated the combination of STAT3 inhibitors (CRYP, stattic) with SBE in Akata-BX1 cells, which led to the more reduction of cell viability (**Fig 6G**) as well as the increased level of EBV ZEBRA protein (**Fig 6H**). In addition, combination of CRYP or stattic with SBE even further enhanced the hIgG-induced EBV lytic reactivation in Akata-BX1 cells **(Fig S6B)**.

**Figure 6.**
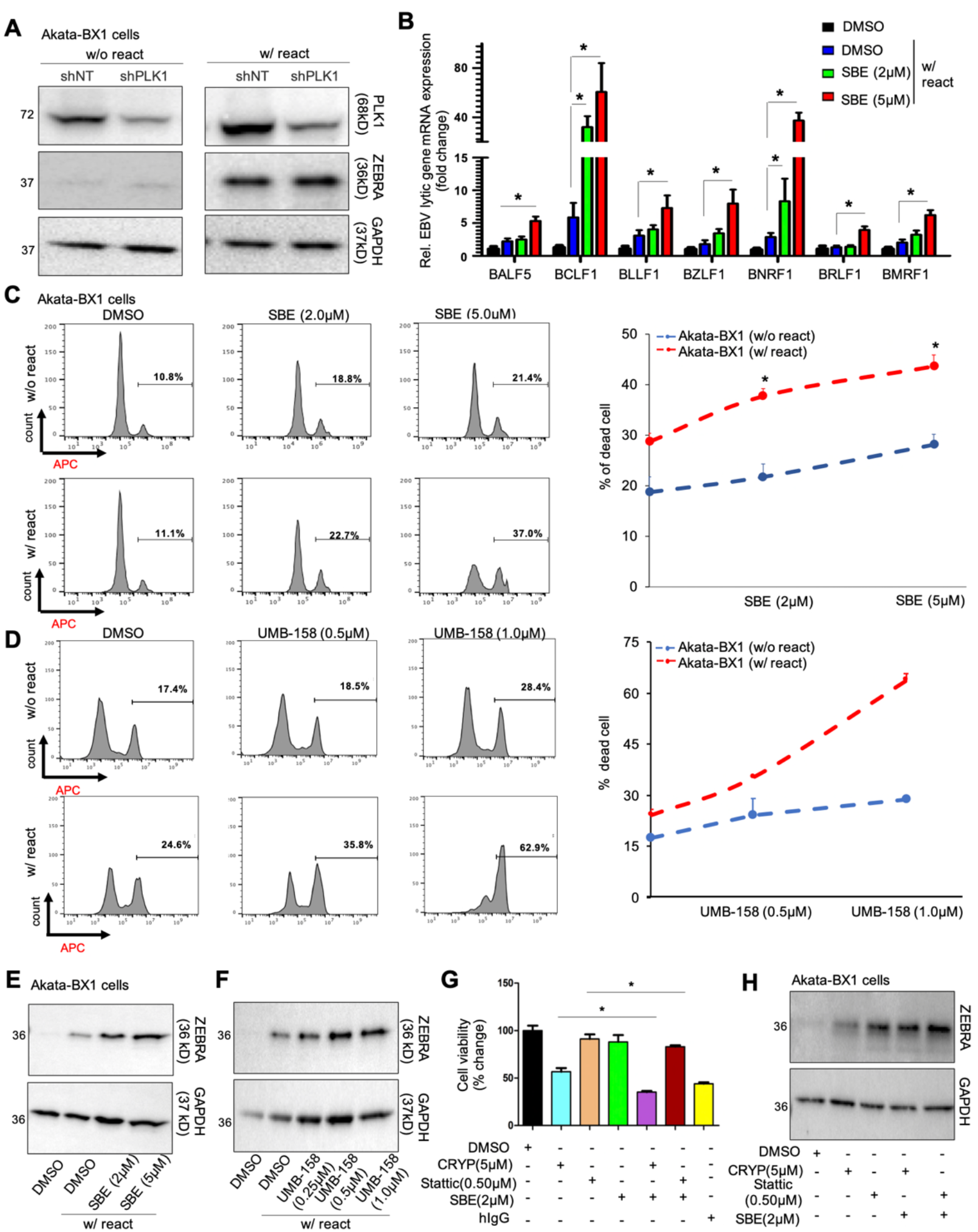
Inhibition of PLK1 benefits EBV reactivation and promotes death of EBV-reactivated tumor cells. (A) Akata-BX1 cells were transiently transfected with the pINDUCER10 vector expressing indicated shRNAs (shNT or shPLK1) using TurboFect™ transfection reagent, and treated with Dox to induce shRNA expression as well as human IgG (hIgG) to induce EBV reactivation. Protein level of PLK1 and EBV lytic gene ZEBRA was measured by immunoblotting. GAPDH was used as a loading control. (B) mRNA level of indicated EBV lytic genes was measured by RT-qPCR and normalized to GAPDH. (C, D) Akata-BX1 cells were treated with SBE (C), UMB-158 (D), or DMSO, in the presence or absence of hIgG. Percentage of dead cells was measured by LIVE/DEAD staining followed by flow cytometry analysis. (E, F) Protein level of EBV ZEBRA in above cells (C, D) was measured by immunoblotting. GAPDH was used as a loading control. (G) Akata-BX1 cells were treated with CRYP or stattic in the presence or absence of SBE. hIgG treatment was used as a positive control to reactivate latent EBV. Cell viability of these cells was measured by ATP-based assays and normalized to DMSO. (H) Protein level of EBV ZEBRA in above cells (G) was measured by immunoblotting. GAPDH was used as a loading control. Results were calculated from n=3 independent experiments and presented as mean ± SD (* p<0.05; two-tailed paired Student t-test).

### PLK1 expression in B cells is elevated in the context of HIV infection

KSHV infection is more common among people living with HIV (PLWH) than in the general population in the world^46,47,48^. As the incidence of B-cell lymphoma among HIV-infected subjects greatly exceeds that of the general population^49^, we compared the PLK1 expression in B cell lymphoma samples from both HIV-negative and positive subjects. PLK1 mRNA abundance were found to be markedly elevated in HIV-positive lymphoma samples compared to HIV-negative samples (**Fig 7A**). This observation led us to specifically investigate the role of PLK1 expression in B cells between HIV-negative and positive subjects. B cells were isolated from peripheral blood mononuclear cells (PBMCs) of these donors (**Fig 7B**), and PLK1 mRNA and protein levels were measured by qPCR and immunostaining respectively. Increased PLK1 mRNA level **(Fig 7C)** was consistent across HIV+ subjects compared to healthy donors. Increased PLK1 protein level was also observed in HIV+ subjects (**Fig 7D, E**). We further determined the impact of HIV infection on PLK1 expression in B cells *in vitro*. Jurkat cells were subjected to HIV-1 IIIB infection, verified by immunoblotting of HIV-1 Gag p24 protein **(Fig S7A)**. HIV-infected Jurkat cells were co-cultured with CFSE-stained BJAB cells, which led to the moderate increase of PLK1 protein level measured by IFAs once compared to BJAB cells co-cultured with non-infected Jurkat cells (**Fig 7F**).

**Figure 7.**
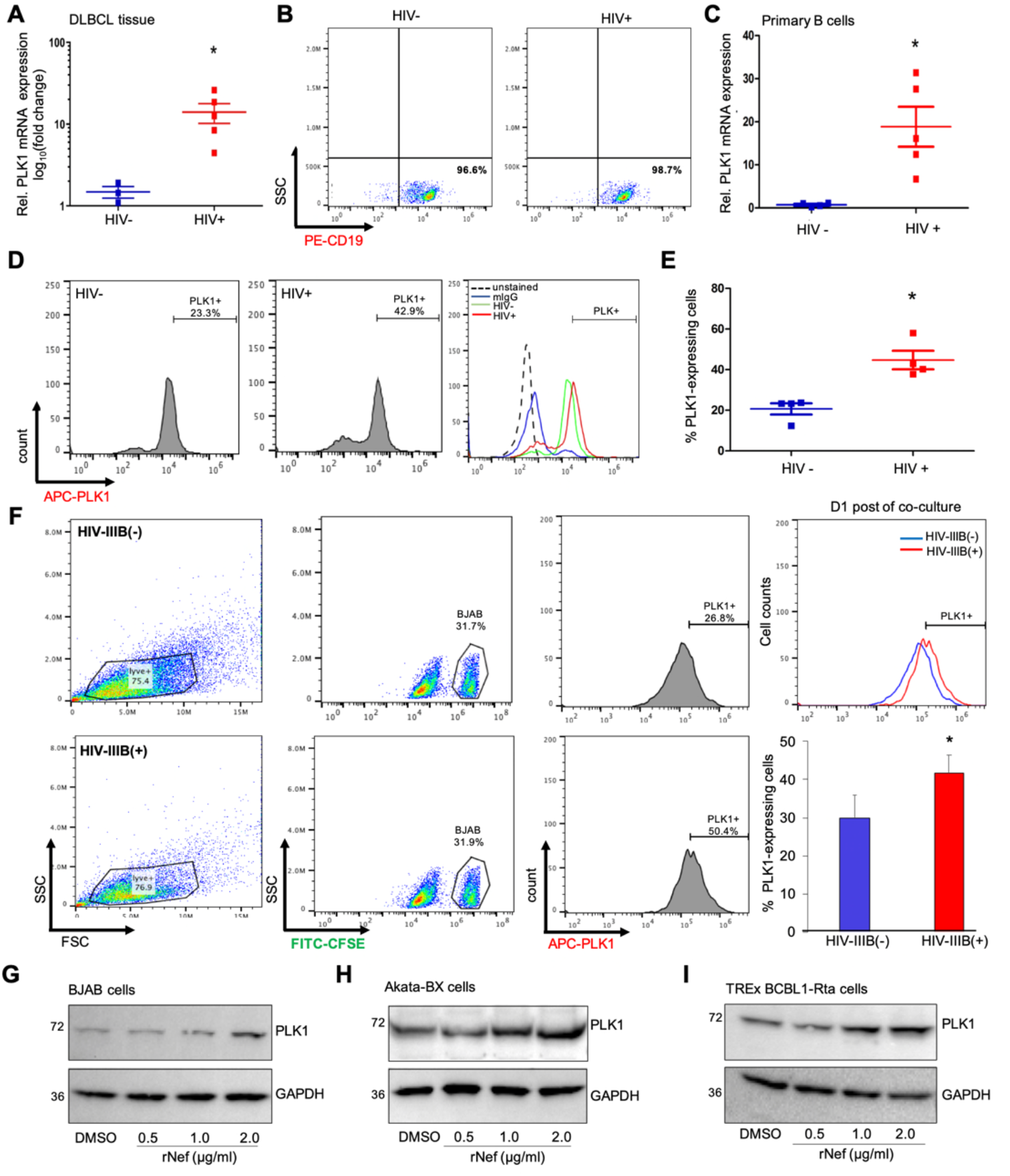
PLK1 expression is induced in B cells in the context of HIV infection. (A) Total RNAs were extracted from the fixed diffuse large B-cell lymphoma (DLBCL) tissue samples of HIV-negative (HIV-, n=3) and HIV-positive (HIV+, n=5) individuals. mRNA level of PLK1 in these tissue samples was measured by RT-PCR and normalized to GAPDH. (B) A PE-conjugated CD19 antibody was used for B cell enrichment from PBMCs of HIV-infected, aviremic patients (HIV+, n=5) or healthy donors (HIV-, n=5). Purity of isolated CD19^+^ B cells was confirmed by flow cytometry analysis. (C) mRNA level of PLK1 in above isolated B cells (B) was measured by RT-PCR and normalized to GAPDH. (D) Protein level of intracellular PLK1 in above isolated B cells (B) was measured by immunofluorescence. Percentage of PLK1-expressing cells was determined by flow cytometry analysis. (F) HIV-1 IIIB infected Jurkat cells were co-cultured with BJAB cells that were pre-stained with CFSE. Protein level of intracellular PLK1 in CFSE-labeled BJAB cells was measured by immunofluorescence. Percentage of PLK1-expressing cells was determined by flow cytometry analysis. (G-I) Protein level of PLK1 in BJAB (G), Atata-BX1 (H), or TREx BCBL1-Rta (I) cells treated with recombinant Nef (rNef) protein at the increasing dose or DMSO was measured by immunoblotting. GAPDH was used as a loading control. Results were calculated from n=3 independent experiments and presented as mean ± SD (* p<0.05; two-tailed paired Student t-test).

We previously showed that HIV Nef protein contributes to the upregulation of PLK1 in HIV-infected T cells^20^. Nef is a ∼27 kDa myristoylated protein encoded by the highly variable *nef* gene located at the 3’ end of HIV genome. Following HIV infection or upon HIV viral reactivation, Nef is expressed as one of the earliest and most abundant viral proteins^50,51^. Nef can be secreted from infected or transfected cells, and detected in the plasma of HIV-infected individuals with undetectable plasma HIV RNA despite of cART suppression^52,53,54,55^. It is generally believed that peripheral Nef protein is produced from HIV-infected cells located in lymphoid tissues as a major HIV reservoir^56,57^. Recently, it has been reported that HIV Nef protein promotes latent infection of KSHV^58^, which was confirmed in our assays that treatment of recombinant Nef (rNef) protein suppresses KSHV lytic reactivation in Dox-treated TREx BCBL1-Rta cells without any cytotoxicity (**Fig S7B,C**). Such function of Nef is in line with PLK1’s role in regulating KSHV latency. Thus, we expected that Nef is the HIV viral protein that mediates the PLK1 induction in B cells. BJAB cells were treated with rNef protein at the increasing dose, which indeed resulted in the upregulation of PLK1 protein **(Fig 7G)**. Similar effect was observed in Akata-BX1 **(Fig 7H)** and TREx BCBL1-Rta **(Fig 7I)** cells treated with rNef protein.

### Inhibition of PLK1 reduces viral reservoirs of KSHV and EBV in B cells

As PLK1 inhibition facilitates lytic switch of KSHV/EBV latency and promotes cell death of KSHV/EBV-infected B lymphoma cells undergoing viral reactivation, we expected that use of PLK1 inhibitors facilitates the reduction or elimination of KSHV/EBV viral reservoirs in these cells. To test this, we prepared a serial dilution of BCBL1 (KSHV+) or Akata-BX1 (EBV+) cells within the KSHV/EBV-negative BJAB cells. BCBL1:BJAB (**Fig 8A**) and Akata-BX1:BJAB (**Fig 8B**) cell mixtures were treated with TPA/NaB and hIgG respectively to induce viral reactivation in the presence or absence of SBE. It was evident that addition of SBE to KSHV/EBV LRAs leads to the significant reduction of KSHV/EBV viral reservoirs in B cells through measurement of viral DNA copy number by RT-qPCR. Surprisingly, treatment of SBE alone also showed the similar effect in these cell mixtures (**Fig S8**), likely due to that although it is weak PLK1 inhibition can still induce certain level of viral reactivation of KSHV and EBV in these cells.

**Figure 8.**
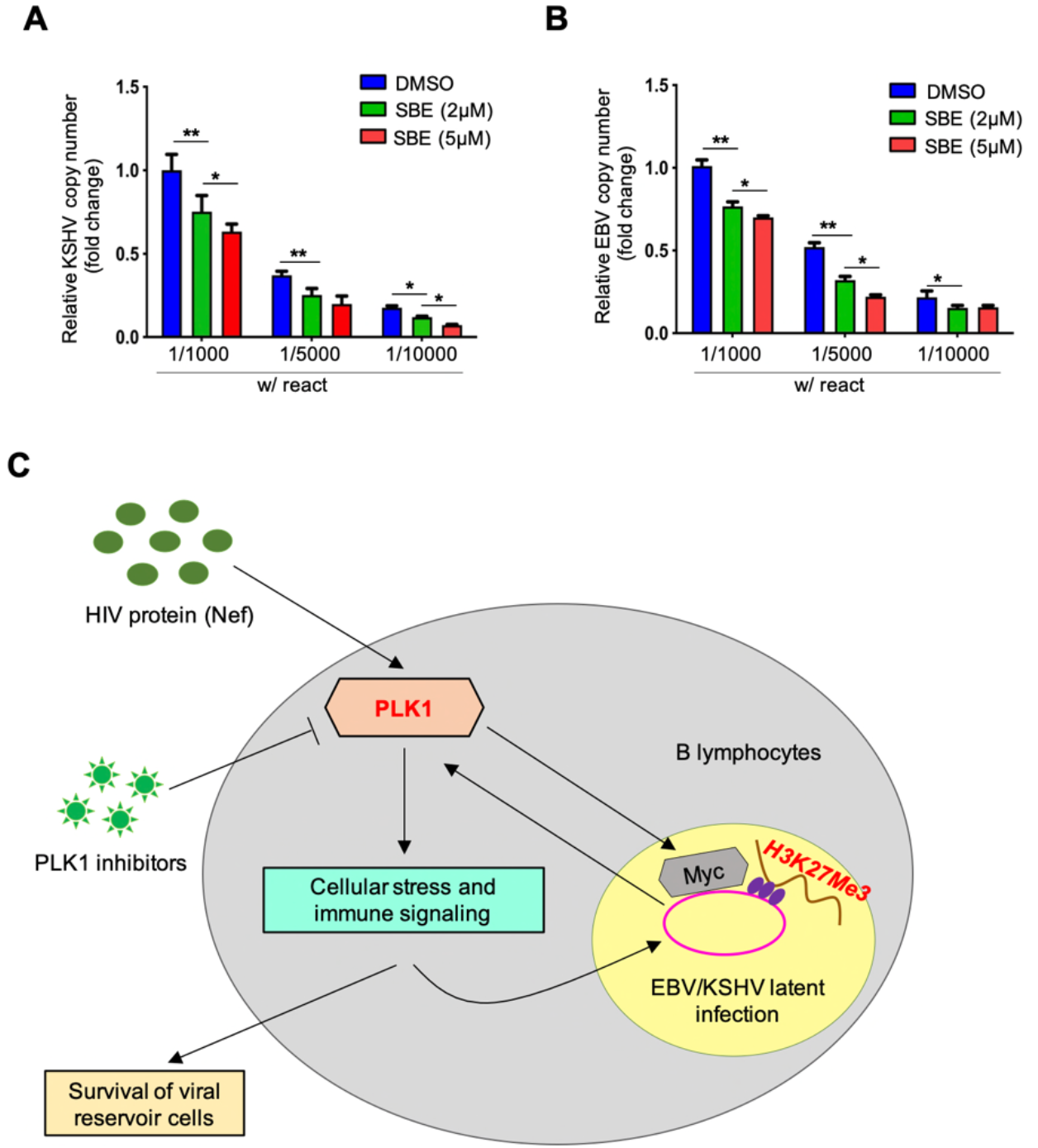
Inhibition of PLK1 reduces KSHV/EBV viral reservoirs in B cells. (A, B) A serial dilution of TREx BCBL1-Rta (A) or Akata-BX1 (B) cells within KSHV/EBV-negative BJAB cells was prepared, followed by treatment of Dox or hIgG respectively to induce viral reactivation, in the presence or absence of SBE. Copy number of viral DNA genome was measured by qPCR using primers that target ORF73/LANA (KSHV) or EBNA1 (EBV) respectively, and normalized to GAPDH. Results were calculated from n=3 independent experiments and presented as mean ± SD (* p<0.05; two-tailed paired Student t-test). (C) A schematic model illustrating the roles of PLK1 in regulating viral latency of human gamma-herpesviruses. PLK1 expression is induced by KSHV infection or in the scenario of HIV infection. PLK1 upregulation would not only support the maintenance of KSHV latency via stabilization of c-Myc and thus increase of H3K27me3 level, but also boost survival of KSHV-infected tumor cells via PLK1’s known multifaceted roles in regulating cellular stress and immune signaling. Hence, PLK1 inhibitors would facilitate lytic switch of KSHV latency and promote death of KSHV-reactivated tumor cells, overall inducing an efficient viral oncolysis and resulting in “killing” of KSHV-infected tumor cells as well as reduction of KSHV viral reservoirs. PLK1 may play such similar roles in regulation of EBV viral latency as well.

## Discussion

Our results showed that both KSHV *de novo* infection and lytic switch from latency lead to the induction of PLK1 (**Fig 1A, B**). Our earlier studies also found that HIV infection induces PLK1 in a PI3K-depdenent manner^20^. It has been reported that KSHV *de novo* infection indeed activates PI3K and that several KSHV proteins, including K1, viral G protein-coupled receptor (vGPCR), vIL-6, and ORF45, participates in it^59^. These KSHV viral proteins also express in latent and lytic switch phases, which would likely result in PLK1 induction as well. Additionally, we also showed that HIV Nef protein is able to induce PLK1 upregulation in B cells (**Fig 7G-I**). It makes sense since HIV Nef protein is similar as PLK1 and also promotes KSHV latency, indicating a functional linage from Nef to PLK1. Interestingly, it has been reported that Nef indeed activates PI3K through multiple mechanisms^60,61,62^: Nef recruits and activates PI3K through association with the Src family tyrosine kinase (SFK) and the tyrosine kinase ZAP-70^61^; Nef binds to the regulatory subunit (p85) of PI3K and activates the Nef-associated p21-activated kinase (PAK)^63^. It has been reported that KSHV/HIV co-infection strongly correlates with cancer progression^64^. Thus, impacts from direct KSHV infection and secreted HIV Nef protein may coordinate PLK1 induction, benefiting KSHV persistent infection and survival of KSHV-infected B lymphocytes. Although it was not tested in our studies, EBV infection likely causes PLK1 induction as well since it has been shown that certain EBV lytic and latent proteins are capable of PI3K activation^65^.

A key finding of our studies is that PLK1 promotes viral latency of gamma-herpesviruses since its depletion (**Figs 1C-F** and **6A,B**) or inhibition (**Figs 2, S2B**, and **6E**,**F**) facilitates viral lytic switch from latency for both KSHV and EBV. Our further mechanistic investigation unraveled that such inhibitory function of PLK1 is likely through regulation of c-Myc protein and histone methylations, particularly H3K27me3 **(Fig 4**). It has been already known that c-Myc is deregulated in ∼30% of human cancers, including B and T cell lymphomas^34,66^, and that it is required for the maintenance of viral latency of both KSHV^35^ and EBV^67^. More importantly, there are several reports delineating the role of PLK1 in regulation of c-Myc. PLK1 inhibition can result in the destabilization of c-Myc protein^19^ as well as the reduction of c-Myc phosphorylation interfering with its transcriptional activities^68^. It is still not clear how exactly c-Myc contributes to viral latency of KSHV and EBV. However, there are several studies suggesting that c-Myc modulates the histone lysine methyltransferases (EZH2, G9a) that count for histone H3 methylations, H3K9me3 and H3K27me3^69^. Interestingly, our results showed that PLK1 inhibition results in the significant reduction of H3K27me3 at both global and local levels at KSHV lytic loci (**Fig 4**), but not H3K9me3 (**Fig S4B**). Indeed, H3K27me3 inactive mark associates with the repressed transcription of KSHV lytic genes^70^. Above all, our studies demonstrated the previously unreported role of PLK1-c-Myc-H3K27me3 axis in promoting KSHV latency. Although it was not tested in our studies, such axis likely has the similar impact on EBV latency given than c-Myc contributes to EBV latency as recently described^67^.

PLK1 also plays a critical role in supporting cell survival, and hence we noticed that PLK1 inhibition further promotes the death of KSHV/EBV-reactivated B lymphoma cells (**Figs 3A,B, S3**, and **6C,D**). However, we also showed that PLK1 inhibition (**Fig S2A**) yields no obvious effect on cell death of SLK cells harboring latent KSHV although it still facilitated KSHV lytic reactivation in these cells (**Fig 2**). It is likely due to that different types of cells possess different sensitivity to the loss or inhibition of PLK1. Nevertheless, our results clearly demonstrated that PLK1 inhibition would really benefit the viral oncolysis approach to treat KSHV/EBV-positive B lymphomas, especially since KSHV infection would cause PLK1 induction in these cells. PLK1 can contribute to cell survival through multiple mechanisms. As shown by our results (**Fig 3E**), PLK1 inhibition deregulated expression of apoptosis-related genes (BAX, BCL2), indicating that cell apoptosis is one type of cell death induced by PLK1 inhibition. We also showed that PLK1 inhibition causes the destabilization of c-Myc that contributes to the release of cell cycle brakes^71^. Thus, PLK1 inhibition would interfere with cell cycle progression, as our results demonstrated that PLK1 inhibition suppresses the expression of c-Myc-controlled cyclin gene expression (**Fig S4A**). In addition, PLK1 also plays a key role in other cell signaling pathways, such as cellular responses to stresses, including DNA damage^72^ and reactive oxygen species (ROS)^73^, as well as immune signaling pathways, including JAK/STAT^31^, NF-kB^74^, RIG-I/MAVS^75^, and Toll-like receptor (TLR)^76^, which all contribute to both viral latency of KSHV/EBV and cell survival. This aspect will be further investigated in the context of KSHV/EBV infection of tumor cells in future studies.

As we previously noted, depletion or inhibition of PLK1 alone is not sufficient to induce KSHV/EBV lytic reactivation. The experimental reagents, such as Dox, hIgG, and TPA/NaB, are not appliable to actual use in clinic. Thus, we seek for reported reagents that are capable of viral reactivation for KSHV/EBV and also suitable for potential future clinic use. Recently, it has been reported that KSHV infection induces STAT3 phosphorylation^77^ and inhibition of STAT3 causes KSHV reactivation in B lymphoma cells^39^. Some of STAT3 inhibitors are currently under evaluation in clinical trials for treating cancers^40,41^. Our results showed that combination of PLK1 and STAT3 inhibitors (CRYP, Stattic) is promising to efficiently induce not only the lytic reactivation of both KSHV (**Fig 5A,B,D,E**) and EBV (**Fig 6H**) but also the cell death of KSHV/EBV-reactivated B lymphoma cells (**Figs 5C,F** and **6G**). This formulates an attractive regimen to specifically target KSHV/EBV-infected tumor cells, and induce reactivation of latent KSHV/EBV and death of these cells through the viral oncolytic approach, which is worthy of further investigation *in vivo*. This would benefit HIV-infected individuals even more since our earlier results also showed that PLK1 inhibition facilitates the “killing” of HIV-infected CD4^+^ T cells and leads to the reduction of HIV viral reservoirs^20^. Although the antiretroviral therapy is effective to control HIV infection^78^, the complications associated with KSHV co-infection remains a significant problem. There was clear evidence of poor overall survival of primary effusion lymphoma (PEL). Recently, one study showed that only 10.2 months of survival for HIV-infected PEL patients receiving multi-drug anticancer treatment^79^. Thus, the regimen of PLK1 and STAT3 inhibitors will be valuable to eliminate KSHV/EBV-infected tumor cells as well viral reservoirs of gamma-herpesviruses (KSHV, EBV) and HIV in these patients.

As summarized in **Fig 8C**, our studies identified that PLK1 is induced by infection of gamma-herpesviruses (KSHV, EBV) or in the context of HIV infection, which in return promotes KSHV/EBV viral latency as well as survival of virus-infected tumor cells. PLK1 inhibition would destabilize c-Myc and reduce histone H3 methylation (H3K27me3), not only facilitating viral reactivation but also promoting death of KSHV/EBV-reactivated tumor cells. Other PLK1-participating cellular stress and immune signaling pathways may also be involved. Our studies also proposed a combinatory regimen of PLK1 and STAT3 inhibitors for the efficient reactivation of latent KSHV/EBV as well as induction of cell death of KSHV/EBV-infected tumor cells, leading to the significant reduction of KSHV/WBV viral reservoirs.

## Acknowledgement

We would like to thank Dr. Prashant Desai for HEK293.r219 cells and Dr. Renfeng Li for Akata-BX1 cells. We would also like to thank Dr. Jae Jung for TREx BCBL1-Rta cells and Dr. SJ Gao for iSLK.r219 and iSLK.BAC16 cells. UMB-158 compound was kindly provided by Dr. Wei Zhang from University of Massachusetts at Boston. This work was supported by grants to N.S. [R03DE029716] and J.Z. [R01DE025447, R01AI150448] from the National Institute of Health.

## Materials and Methods

### Cells

Jurkat Clone E6-1 (Cat. #177) and SLK cells were received from the NIH AIDS reagent program. TREx BCBL1-Rta cells^80^ were maintained in RPMI 1640 medium (Invitrogen) supplemented with 10% fetal bovine serum (FBS; Invitrogen), 1% Pen-Strep, 20 mg/ml hygromycin B and 2 mM L-glutamine (Invitrogen). KSHV-positive BCBL1 cells were grown in RPMI 1640 supplemented with 20% heat-inactivated FBS and 2 mM l-glutamine^80^. BJAB cells were maintained in RPMI 1640 medium (Invitrogen) supplemented with 10% fetal bovine serum (FBS; Invitrogen) and 1% Pen-Strep. iSLK.r219 cells^81^ were maintained in DMEM medium (Corning) containing 10% FBS, 1% Pen-Strep, 10 μg/ml puromycin (Corning), 250 mg/ml Geneticin (Corning), and 400 mg/ml hygromycin B (Corning). iSLK.BAC16 cells were maintained in the presence of 1 μg/ml puromycin, 250 μg/ml G418, and 1,200 μg/ml hygromycin B^23^. Akata-BX1 cells were maintained in RPMI 1640 supplemented with 10% FBS. EBV-negative Akata cell line was clonally selected for the loss of viral episome from the original EBV-positive Akata Burkitt’s Lymphoma cell line, which was subsequently re-infected with EBV BX1 strain and selected with neomycin^82^.

### Compounds

DMSO was purchased from Fisher Scientific. SBE 13 HCl were purchased from Selleck Chemicals. Recombinant HIV Nef protein (rNef) were provided by NIH AIDS reagent program. The UMB-158 PLK1/BET dual inhibitor was synthesized according to an earlier publication^28^ and kindly provided by Wei Zhang (University of Massachusetts at Boston). The STAT3 inhibitors, cryptotansinone and stattic, were purchased from Sigma-Aldrich. 12-O-Tetradecanoylphorbol 13-acetate (TPA, Cat#P8139) and sodium butyrate (NaB, Cat#AAA1107906) were purchased from Sigma-Aldrich and Fisher Scientific, respectively. Doxycycline (Dox) was obtained from Fisher Scientific (Cat#BP2653-1). Human IgG (Cat#55087) were purchased from MP biomedicals.

### Viruses

iSLK.BAC16 cells were treated with 1µg/mL Dox and 1mM NaB for 48h. Supernatants were collected 2 dpi, centrifuged (400x g) for 10 mins to remove cellular debris, and filtered through the 0.45µm filter. The harvested supernatants containing KSHV.BAC16 viruses were incubated with pre-seeded SLK cells cultured in the 1:1 ratio of fresh media along with 8µg/mL polybrene via spinoculation (2500 rpm) for ∼ 4h at 37°C. HIV-1 IIIB wild-type virus was kindly provided by the NIH AIDS reagent program^83,84^. Jurkat cells were subjected to HIV-1 IIIB infection at multiplicity of infection (MOI) = 1 for 5 days^20,84^.

### Antibodies

ChIP-grade mouse anti-PLK1 (Cat. # SAB1404220) and mouse IgG (Cat. #sc2025) antibodies were obtained from Sigma Aldrich. Anti-FLAG (Cat. #2368) antibody was purchased from Cell Signaling Technology. Mouse anti-V5 (Cat. # R960-25), anti-HA (Cat. # 26183), HRP-conjugated goat anti-Mouse IgG (H+L) secondary antibody (Cat # 31430), HRP-conjugated goat anti-Rabbit IgG (H+L) secondary antibody (Cat # 31460) were purchased from Thermo Fisher Scientific. Antibodies against c-Myc (Cat. # 9E10: sc-40), HHV-8 K8.1A/B (Cat. # sc-65446), EBV ZEBRA (Cat. # sc-53904), GAPDH (Cat. # sc-32233) were purchased from Santa Cruz Biotechnology. Antibodies against H3K27Me3 (Cat. # 39155), H3K9Me3 (Cat. # 61013) were purchased from Active Motif, and histone H3 antibody (Cat. # 9715S) was purchased from Cell Signaling Technology.

### siRNAs and shRNAs

For siRNA knockdown assays, PLK1 siRNA (s449) or non-targeting control (NT) siRNA (AM4636) was purchased from Thermo Fisher Scientific. TREx BCBL1-Rta cells were transiently reverse-transfected with siPLK1 (50nM) or siNT (50nM) using RNAiMAX reagents^85^ according to the manufacturer’s instruction. Cells were kept in culture for 72 hrs. These cells were treated with Dox (2μg/ml) to induce KSHV reactivation. For shRNA assays, endogenous PLK1 was knocked down in Akata-BX1 cells by using its shRNA expressed from the pINDUCER10 Dox-inducible lentiviral vector, according to the reported protocol^86,20^. Briefly, pINDUCER10 expressing shRNAs (**Table S1**) was transiently transfected in Akata-BX1 cells using TurboFect™ transfection reagent (Thermo fisher scientific) according to the manufactures protocol. Cells were kept in culture for 72 hrs. These cells were treated with Dox to induce shRNA expression and human IgG (2 µg/ml) to induce EBV reactivation for 48 hrs.

### Isolation of primary B cells

Peripheral blood mononuclear cells (PBMCs) from HIV-infected individuals were acquired from Vitrologic Biological source (Charleston, SC). PBMCs from healthy donors were purchased from STEMCELL™ Technologies (Cambridge, MA). PBMCs from HIV+/- donors were subjected to B-cell isolation. Briefly, PBMCs were captured by using an anti-CD19 antibody conjugated to colloidal paramagnetic microbeads (B-cell isolation kit; Miltenyi Biotec, Bergisch-Gladbach, Germany) and passed through a magnetic separation column (LS; Miltenyi Biotec). The purity of isolated B cells was over 95% as assessed by flow cytometry analysis of PE-CD19^+^ cells (Miltenyi Biotec) ^87,88^.

### Chromatin immunoprecipitation (ChIP)

ChIP assay was conducted according to the manufacturer protocol (Millipore Sigma, cat # 17-395) as previously described^89,90^. Briefly, TREx BCBL1-Rta cells were cross-linked by using 0.5% formaldehyde, followed by treating with 1x glycine to quench the reaction. Cells were washed with cold 1x PBS, and nuclei were isolated using nuclei isolation buffer with rigorous vertexing. Nuclear lysates were sonicated for 2 mins to fragment genomic DNAs. All extraction procedure was carried out in presence of protease inhibitor cocktail. 1% input were separated before the next step. The lysates were incubated with pre-washed Magna ChIP A/G beads along with specific antibodies or control IgG for overnight at 4°C. Samples were subjected to the subsequent washes and eluted through magnetic separator. Each sample was treated with proteinase K. To reverse the cross-linking, the eluted samples were incubated at 65°C for 2 hrs and then 95°C for 15 mins. Next, final magnetic separation was performed to elute the samples. KSHV lytic gene expression was quantified by real-time PCR. Input (1%) was used for qPCR analysis.

### Confocal and fluorescence microscopy

For iSLK.r219 cells, fluorescence was measured on a BioTeK plate reader by using its GFP and RFP channels. Expression of GFP and RFP proteins from KSHV.r219 viral strain are respectively driven by the eIF1a promoter and KSHV lytic PAN promoter^91^. SLK and iSLK.BAC16 cells were seeded onto the high precision cover glasses (Bioscience Tools, Cat#CSHP-No1.5-13) pre-coated with poly-L-lysine (R&D Systems, Cat#3438-100-01) for 45 mins in a 24-well plate. These cells were induced with 1µg/mL Dox for 48 hrs. The KSHV.BAC16 *de novo* infected SLK cells were also collected in coverslips. Cells were rinsed and fixed with 4% paraformaldehyde at room temperature (RT), followed by permeabilization with 0.05% triton-X reagent for 10 mins at RT. Cells were washed with 1x PBS and incubated with 5% FBS for 1 hr at RT. Cells were further incubated with an anti-mouse PLK1 antibody in 2.5% FBS for overnight in a moist chamber at 4°C. Cells were washed to remove the primary antibody and incubated with the alexa 647-mouse secondary antibody for 1 hr at RT. Cells were stained with Hoechst (Invitrogen) as per manufacturer’s guidelines. Coverslips were rinsed and mounted on slides by using ProLong Glass Antifade Mountant (Invitrogen, Cat#P36982). Slides were left to cure in dark for 24 hrs at RT per manufacturer’s recommendations. Confocal images were acquired by using the ZEISS LSM 700 Upright laser scanning confocal microscope and ZEN imaging software (ZEISS). PLK1 were imaged via TAMRA channel while GFP is constitutively expressed in cells infected with KSHV BAC16 ^23^.

### Immunofluorescence assay (IFA)

Isolated B cells were isolated from HIV+/- subjects. Portion of isolated B cells was incubated with CD19 antibody (1/200 of stock, Milteny) for 30 mins to determine the purity using BD Accuri C6 Plus with corresponding optical filters. The rest of B cells was washed and fixed with 4% paraformaldehyde at RT for 20 mins. Pelleted cells were washed and permeabilized with saponin-containing 1× Perm/Wash buffer (BD Biosciences) as described^20^. Cells were incubated with an anti-PLK1 antibody (200μg/ml) diluted in 1× Perm/Wash buffer for overnight at 4°C, followed by incubation with the fluorophore-conjugated secondary antibody for 1 hr at RT in the dark. The staining buffer (1× D-PBS with 2% bovine serum albumin [BSA]) was added to resuspend the cells, followed by flow cytometry analysis using the BD Accuri C6 Plus with corresponding optical filters. For the Jurkat and BJAB co-culture assays, BJAB cells were labeled separately with CFSE (Thermo Fisher) for 30 mins according to instruction and incubated with HIV-1 IIIB infected Jurkat cells. These cells were incubated with an anti-PLK1 antibody, and subsequently the fluorophore-conjugated secondary antibody. The mean fluorescence intensity (MFI) and the percentage of fluorescence-positive cells were determined by using the FlowJo V10 software.

### Immunoblotting assay

Immunoblotting was carried out using the existing protocol^2084^. In brief, cell pellets were homogenized in ice with 1x RIPA containing protease inhibitor cocktail. The cell lysate was cleared by centrifugation at 12,000 rpm for 10 mins. The protein concentration was measured by BCA kit (Thermo Fisher Scientific). The same amount of protein samples was boiled in 2x SDS loading buffer, separated by SDS-PAGE, and transferred to PVDF membrane. The blots were blocked with 5% skimmed milk in 1x PBS and probed with the specific primary antibodies followed by HRP-conjugated secondary antibodies. Protein bands were visualized with ECL Plus chemiluminescence reagent.

### Cell viability and death assays

The cytotoxicity of compounds was determined by using the ATP-based CellTiter-Glo Luminescent Cell Viability Assay (Promega) and analyzed by the Cytation 5 multimode reader (luminescent mode). Death of virus-infected or compound-treated cells was determined by using the LIVE/DEAD Fixable Far Red Dead Cell Stain Kit (Invitrogen) and analyzed by the BD Accuri C6 Plus (flow cytometry)^20^.

### Cell apoptosis assay

The Vybrant Apoptosis Assay Kit (Thermo Fisher, cat # V13243) was used to measure cell apoptosis. YO-PRO-1 and propidium iodide (PI) nucleic acid staining was performed following the manufactures instructions. YO-PRO-1 selectively passes through the plasma membrane of apoptotic cells and labels them with green fluorescence. Necrotic cells are stained witht the red-fluorescent PI^92^.

### Real-time qPCR (RT-qPCR)

Extraction of genomic DNA was performed by using the DNeasy Blood & Tissue Kit (Qiagen, cat # 69504) according to the manufacture’s instruction. Total RNAs were extracted from the assayed cells by using the RNeasy kit (Qiagen), and 0.2-1 µg of RNA was reversely transcribed using the iScript™ cDNA Synthesis Kit (Bio-Rad). Real-time PCR assay was conducted using the SYBR Premix ExTaq II (Bio-Rad) and gene-specific primers (**Table S1**). The PCR reactions were performed on a Bio-Rad CFX connect qPCR machine under the following conditions: 95 °C for 10 mins, 40 cycles of 95 °C for 15 secs and 60 °C for 1 min. Relative percentage of gene expression was normalized to GAPDH control, and was calculated using the formula: 2 ^(Δ^ ^C^ ^T^ ^of gene−Δ^ ^CT^ ^of^ ^GAPDH)^.

### RNA isolation from paraffin tissue

Paraffin-embedded tissue sections were deparaffinized by using xylene solution twice with each for 5 mins at RT followed by the high-speed centrifugation at 12,000 rpm for 20 mins. Isolated pellets were washed with graded alcohol (100, 90, 70%) and centrifuged at 12,000 rpm for 10 mins. Air-dried pellets were digested with the buffer containing 4M guanidine thiocyanate and 1M Tris-HC1 (pH 7.6) in DEPC-treated water. 0.3% β-mercaptoethanol was added in the digestion buffer in addition to proteinase K (6 mg/mL) for another incubation at 45°C for overnight. 1 vol of phenol-water/chloroform (70/30) was added and vortexed rigorously. The mixture was kept on ice for 15 mins and centrifuged at 12,000 rpm for 20 mins. The upper aqueous phase was collected by avoiding the proteinaceous interface between the two phases, and transferred to a new test tube. 1 vol of isopropanol was added in addition to 1mg/ml glycogen. The aqueous phase was kept at −80°C for 2 hrs, followed by the centrifugation at 12,000 rpm for 20 mins at 4°C. RNA pellets were washed with ethanol (75%) and kept at −20°C for overnight. RNA pellets were centrifuged at 12,000 rpm for 5 mins and air-dried. DEPC-treated water was added to resuspend the pellet and dissolve the RNA.

### Statistical analysis

Statistical analysis was performed in GraphPad PRISM 5 or Excel. Data are presented as mean ± SD of independent experiments (n = 3). * p<0.05 and ** p<0.01 with two-tailed paired Student t-test except specified otherwise.

---

**Figure S1.**
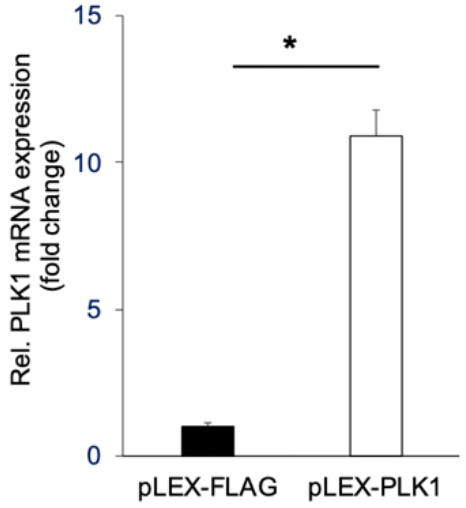
mRNA level of PLK1 in iSLK.r219 cells transiently transfected with the pLEX-FLAG or pLEX-PLK1 vector was measured by RT-qPCR. (* p<0.05; two-tailed paired Student t-test).

**Figure S2.**
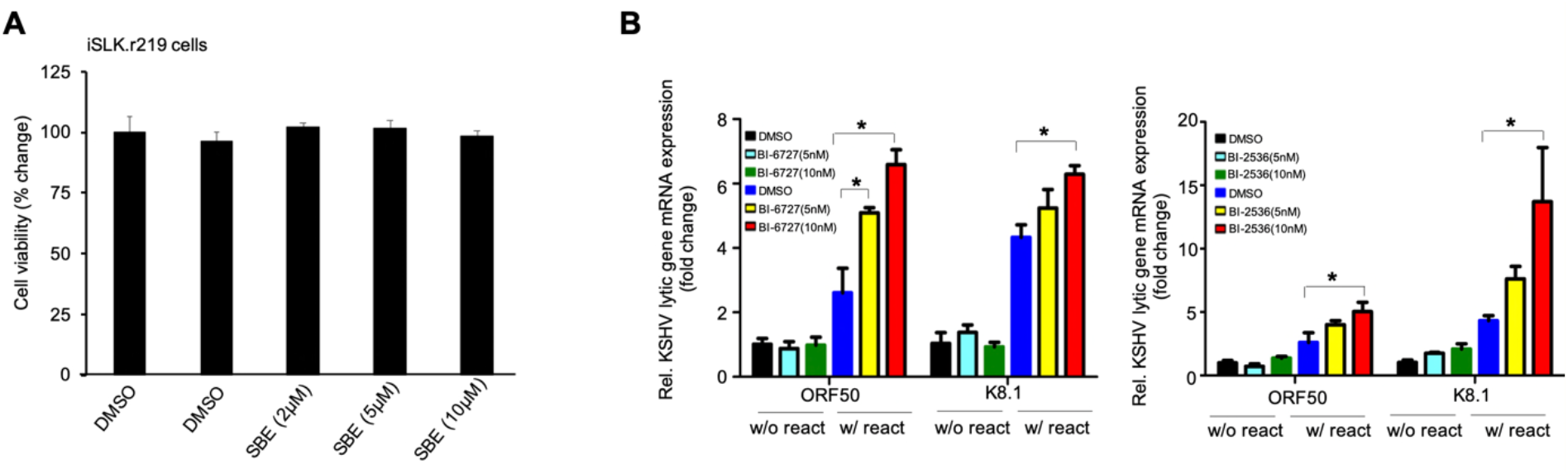
(A) Cell viability of iSLK.r219 cells treated with SBE at the increasing dose as well as Dox was measured by ATP-based assay. (B) mRNA level of KSHV lytic genes (ORF50, K8.1) in TREx BCBL1-Rta cells treated with BI-6727 or BI-2536 in the presence or absence of Dox was analyzed by qPCR and normalized to GAPDH. (* p<0.05; two-tailed paired Student t-test).

**Figure S3.**
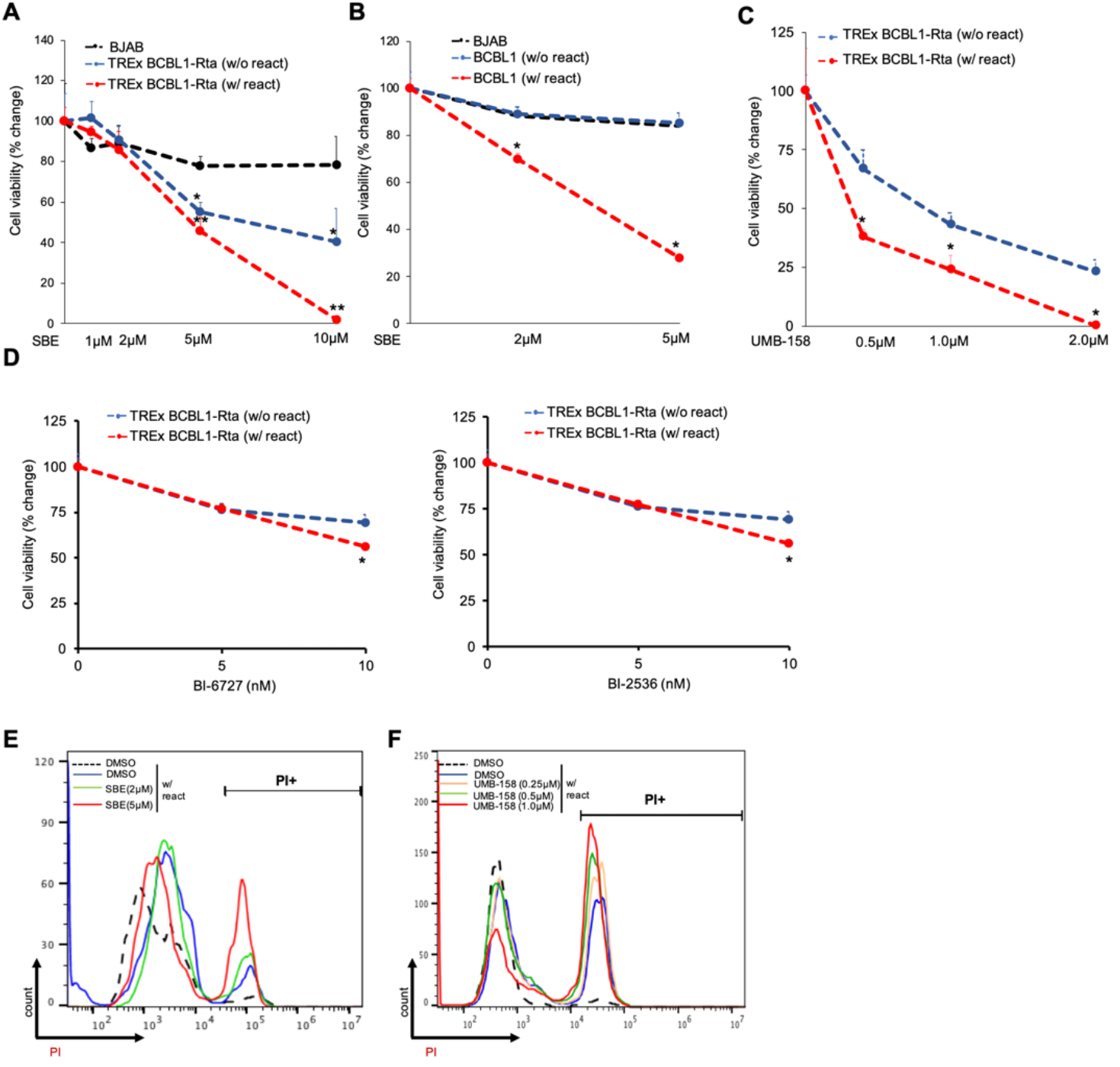
(A, B) Cell viability of TREx BCBL1-Rta (A) or BCBL (B) cells treated with SBE with or without Dox was measured by ATP-based assay. In parallel, cell viability of KSHV/EBV-negative BJAB cells treated with SBE alone was also measured. (C) Cell viability of TREx BCBL1-Rta cells treated with UMB-158 with or without Dox was measured. (* p<0.05, **p<0.001; two-tailed paired Student t-test). (D) Cell viability of TREx BCBL1-Rta cells treated with BI-6727 or BI-2536 in the presence or absence of Dox was measured by ATP-based assay. (* p<0.05; two-tailed paired Student t-test). (E,F) TREx BCBL1-Rta cells were treated with SBE (E), UMB-158 (F), or DMSO, and induced with Dox. Percentage of necrotic cells was analyzed by flow cytometry of PI-stained cells using Vybrant Apoptosis Assay Kit (Thermo Fisher),.

**Figure S4.**
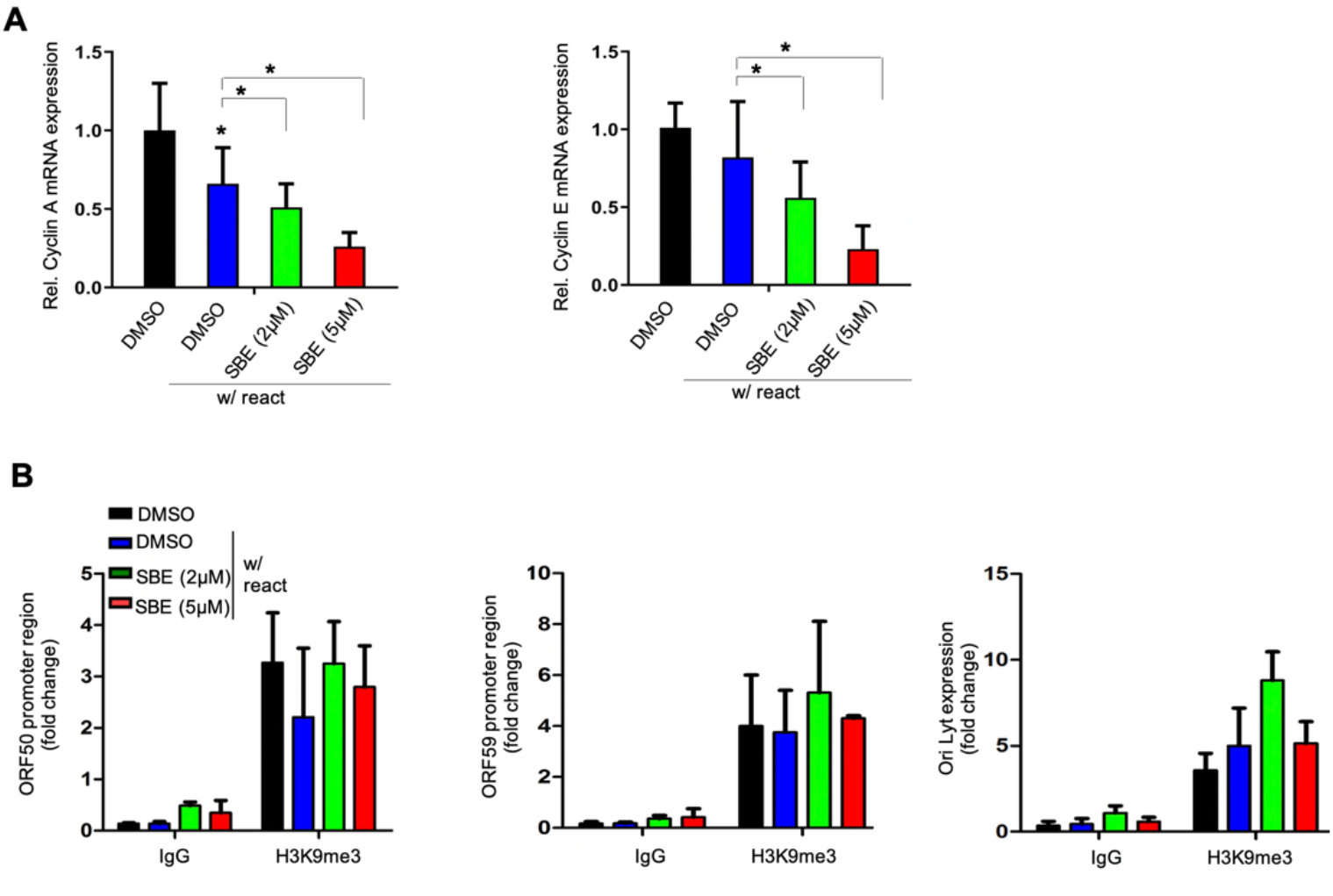
(A) mRNA level of cell-cycle genes, cyclin A and cyclin E, in TREx BCBL1-Rta cells treated with SBE or DMSO in the presence of Dox was measured by RT-qPCR and normalized to GAPDH. (B) TREx BCBL1-Rta cells were treated with SBE or DMSO, and Dox to induce KSHV reactivation. Cell lysates were prepared and subjected to ChIP assay by using antibodies against H3K9me3 or a mouse IgG. Precipitated DNA samples were further analyzed by qPCR by using primers targeting promoter region of KSHV lytic genes (ORF50, ORF59) and OriLyt, and normalized to IgG control. (* p<0.05; two-tailed paired Student t-test).

**Figure S5.**
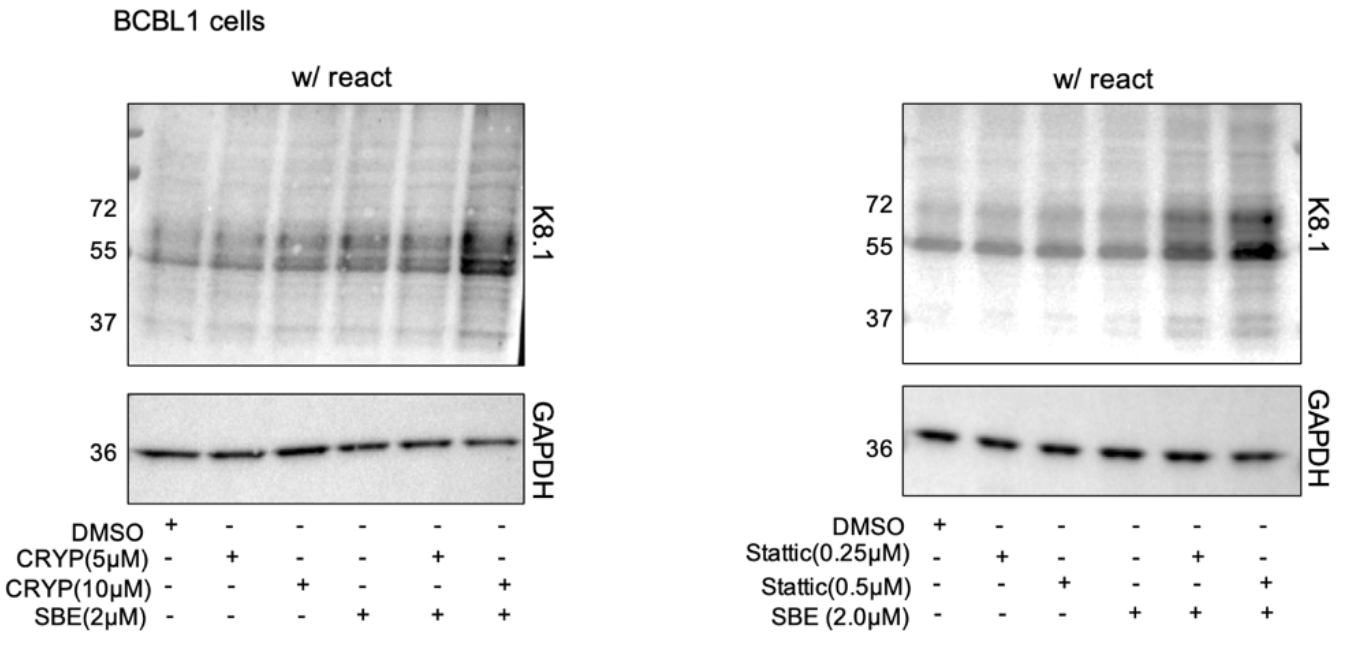
Protein level of KSHV K8.1 in BCBL1 cells treated with CRYP or stattic in the presence or absence of SBE was measured by immunoblotting. GAPDH was used as a loading control.

**Figure S6.**
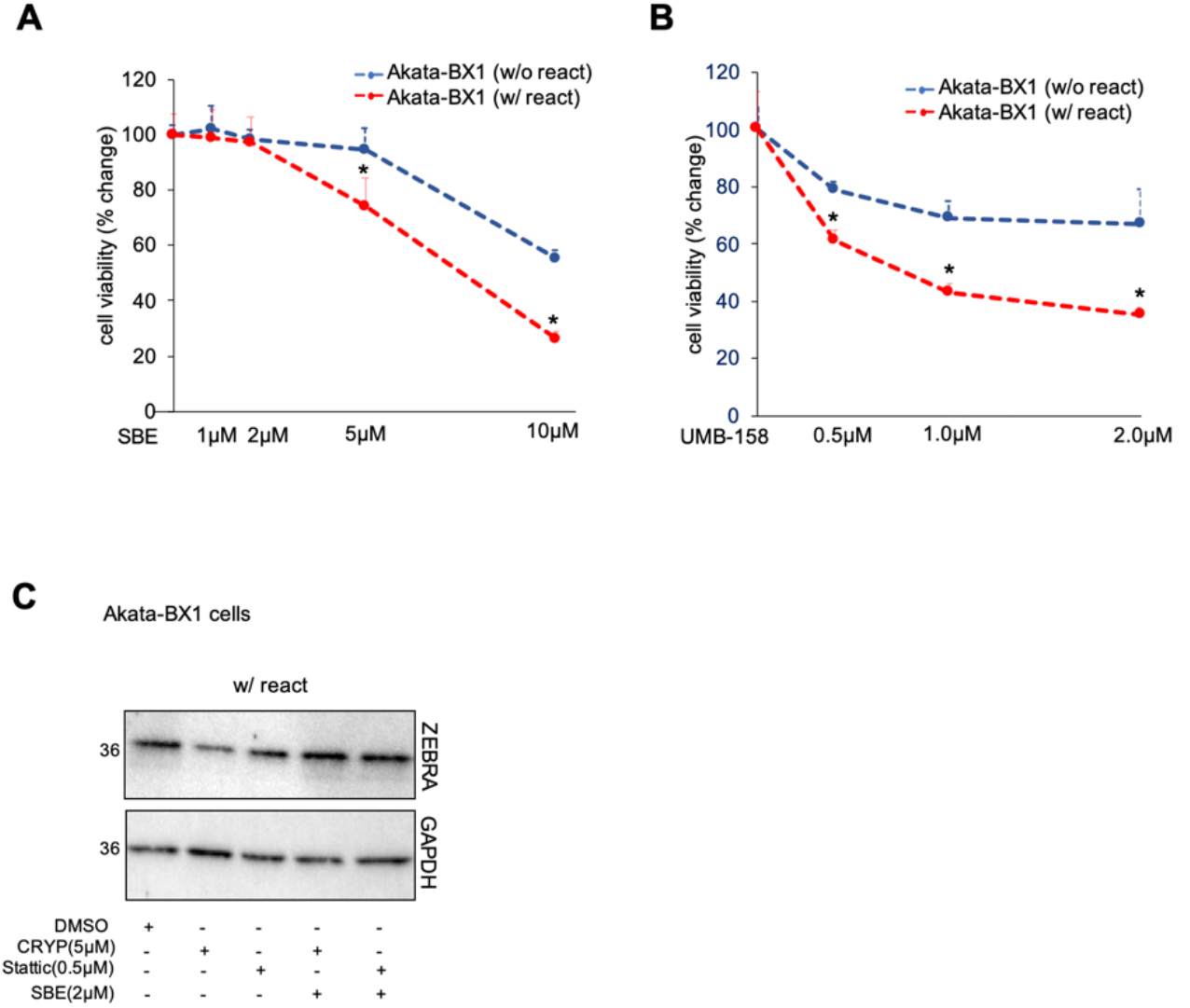
(A) Cell viability of Akata-BX1 cells treated with UMB-158 at the increasing dose with or without hIgG induction was measured by ATP-based assay. (B) Protein level of EBV ZEBRA in Akata-BX1 cells treated with CRYP or stattic in the presence or absence of SBE was measured by immunoblotting. GAPDH was used as a loading control.

**Figure S7.**
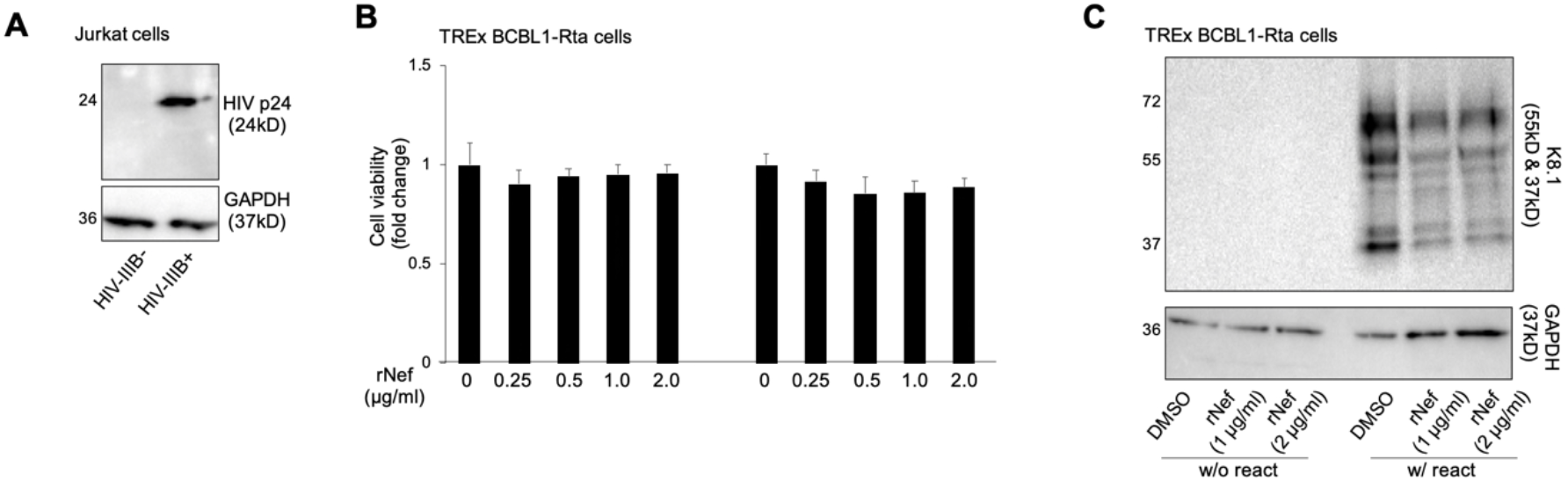
(A) HIV-1 IIIB infection in Jurkat cells was confirmed by immunoblotting of HIV p24 protein. (B) Cell viability of TREx BCBL1-Rta cells treated with rNef protein in the presence or absence of Dox was measured by ATP-based assay. (C) Protein level of KSHV K8.1 in above cells (B) was measured by immunoblotting.

**Figure S8.**
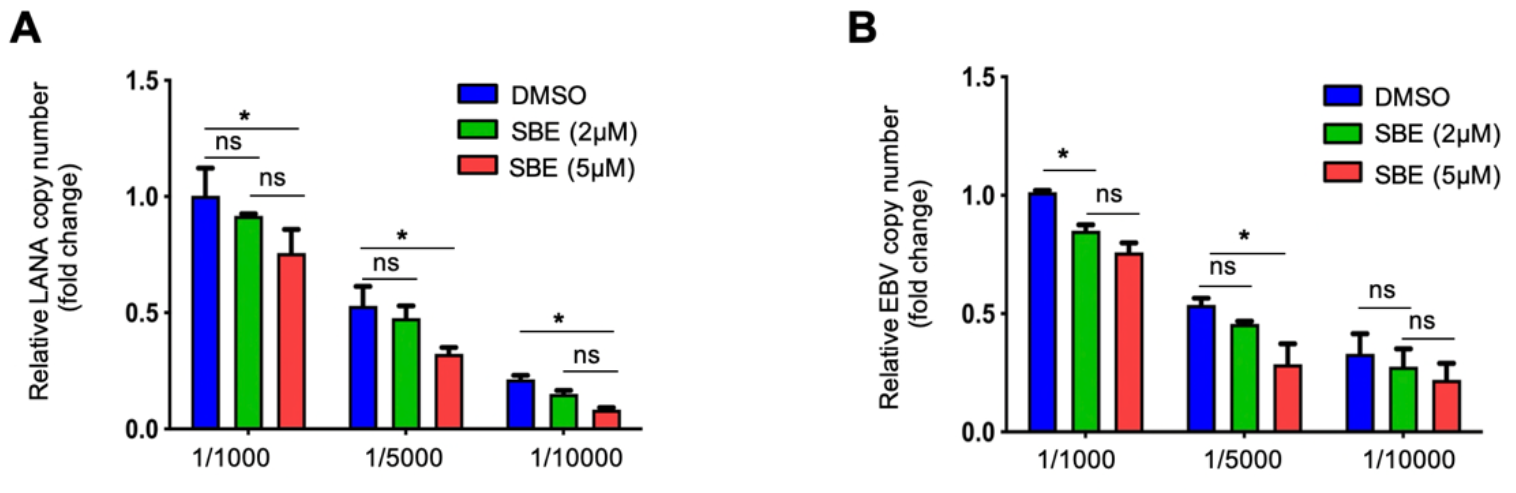
A serial dilution of TREx BCBL1-Rta (A) or Akata-BX1 (B) cells within KSHV/EBV-negative BJAB cells was prepared, followed by treatment of SBE alone or DMSO. Copy number of viral DNA genome was measured by qPCR using primers that target ORF73/LANA (KSHV) or EBNA1 (EBV) respectively, and normalized to GAPDH. (* p<0.05; two-tailed paired Student t-test).

## Notes

### Competing Interest Statement

The authors have declared no competing interest.

### Summary of Updates

We've provided more results presented in new figures.

